# Kinetic modelling of β-cell metabolism reveals control points in the insulin-regulating pyruvate cycling pathways

**DOI:** 10.1101/2021.08.02.454627

**Authors:** Rahul Rahul, Adam R Stinchcombe, Jamie Joseph, Brian Ingalls

## Abstract

Insulin, a key hormone in the regulation of glucose homeostasis, is secreted by pancreatic β-cells in response to elevated glucose levels. Insulin is released in a biphasic manner in response to glucose metabolism in β-cells. The first phase of insulin secretion is triggered by an increase in the ATP:ADP ratio; the second phase occurs in response to both a rise in ATP:ADP as well as other key metabolic signals, including a rise in the NADPH:NADP^+^ ratio. Experimental evidence indicates that pyruvate-cycling pathways play an important role in the elevation of the NADPH:NADP^+^ ratio in response to glucose. In this work we developed a kinetic model for the tricarboxylic acid cycle and pyruvate cycling pathways. We successfully validated our model against recent experimental observations and performed local and global sensitivity analysis to identify key regulatory interactions in the system. The model predicts that the dicarboxylate carrier (DIC) and pyruvate transporter (PYC) are the most important regulators of pyruvate cycling and NADPH production. In contrast, our analysis showed that variation in the pyruvate carboxylase (PC) flux was compensated by a response in the activity of mitochondrial isocitrate dehydrogenase (ICD_m_) resulting in minimal effect on overall pyruvate cycling flux. The model predictions suggest starting points for further experimental investigation, as well as potential drug targets for treatment of type 2 diabetes.

## Introduction

Pancreatic β-cells integrate a variety of nutrient signals to regulate the secretion of the hormone insulin, which plays an important role in the regulation of nutrient homeostasis in adipose, liver and skeletal muscle tissue. The failure of β-cell activity, resulting in dysregulated insulin release, in combination with insulin-sensitive tissues becoming insulin resistant, results in the development of type 2 diabetes [1].

The mechanism by which elevated glucose levels trigger the release of insulin within β-cells has not been fully characterized. Glucose-stimulated insulin-secretion (GSIS) is bi-phasic. The first phase of insulin release is regulated by the K_ATP_-dependent pathway through which increased glucose metabolism mediates a rise in the ratio of adenosine triphosphate (ATP) to adenosine diphosphate (ADP) in the cytosol, causing closure of ATP-dependent potassium (K_ATP_) channels. The resulting depolarization of the cell membrane causes an influx of Ca^2+^, triggering the secretion of insulin vesicles [2]. This first-phase of insulin release occurs within the first ten minutes following glucose stimulation [3]. The second phase of insulin secretion, which is prompted by the K_ATP_-independent pathway (also known as the amplifying pathway), involves an increase in the ATP:ADP ratio along with other metabolic signals, such as a rise in the NADPH:NADP^+^ ratio [3], [4].

The K_ATP_ channel independent pathway relies on metabolism of pyruvate in the mitochondria. In most cell types, pyruvate feeds the tricarboxylic acid (TCA) cycle via pyruvate dehydrogenase (PDH), which generates acetyl-CoA. β-cells are one of the few cell types that express higher levels of the enzyme pyruvate carboxylase (PC), which provides an alternative route for pyruvate entry to the TCA cycle.

When addressing a complex metabolic network like the TCA cycle, it can be difficult to predict the effects of individual genetic or biochemical perturbations on the entire system. Kinetic modelling provides a framework for addressing networks in a systematic and quantitative manner. Analysis of mathematical models aids in the interpretation of experimental data and can guide the design of further experiments to elucidate the underlying biological process. A number of computational models have been developed to describe aspects of the TCA cycle and GSIS [5]–[7]. In this paper, we build on these previous efforts to develop a mathematical model of β-cell metabolism that describes pyruvate cycling pathways.

We describe the development of a detailed model of pyruvate cycling which we validated against published experimental observations of the pyruvate cycling pathways. We successfully fit the model to data from Ronnebaum *et al*. [8] and then validated the model by confirming that it reproduces a wide range of previously published GSIS investigations carried out by ^13^C isotopomer analysis and siRNA mediated knock-down [9]–[16]. We performed local and global sensitivity analysis to identify key control points in the pathway. This analysis reveals that the pyruvate concentration is heavily influenced by the V_max_ of PYC and is robust to perturbations in other system features. Considering features of the pyruvate cycling pathways, the V_max_ values of dicarboxyrate carrier (DIC), cytocolic malic enzyme (ME_c_) and pyruvate carboxylase (PC) have significant influence over the pyruvate level, indicating that these are important control points in TCA cycle anaplerosis. Similarly, the NADPH concentration is heavily influenced by these parameters, establishing a further correlation between pyruvate cycling and GSIS.

## Methods

### Model Development

In most cell types, pyruvate feeds the tricarboxylic acid (TCA) cycle via pyruvate dehydrogenase (PDH), which generates acetyl-CoA. However, β-cells express significant quantities of pyruvate carboxylase (PC), which provides an additional route from pyruvate to the TCA cycle by producing oxaloacetate (OAA_m_) from pyruvate (Figure 1). In β-cells, pyruvate flows into mitochondrial pathways through these enzymes in approximately equal proportion [1]. Pyruvate that enters the TCA cycle via PC can readily be recycled back to pyruvate, either directly from OAA_m_ (via PC) or after further metabolism of OAA_m_ in the TCA cycle, possibly involving both mitochondrial and cytosolic enzyme activity [15]. The experiments of Lu *et al*. revealed that GSIS is related to PC-catalyzed pyruvate cycling, and that PDH-catalyzed conversion of pyruvate to acetyl-CoA does not play a significant role in GSIS [16].

**Figure 1:**
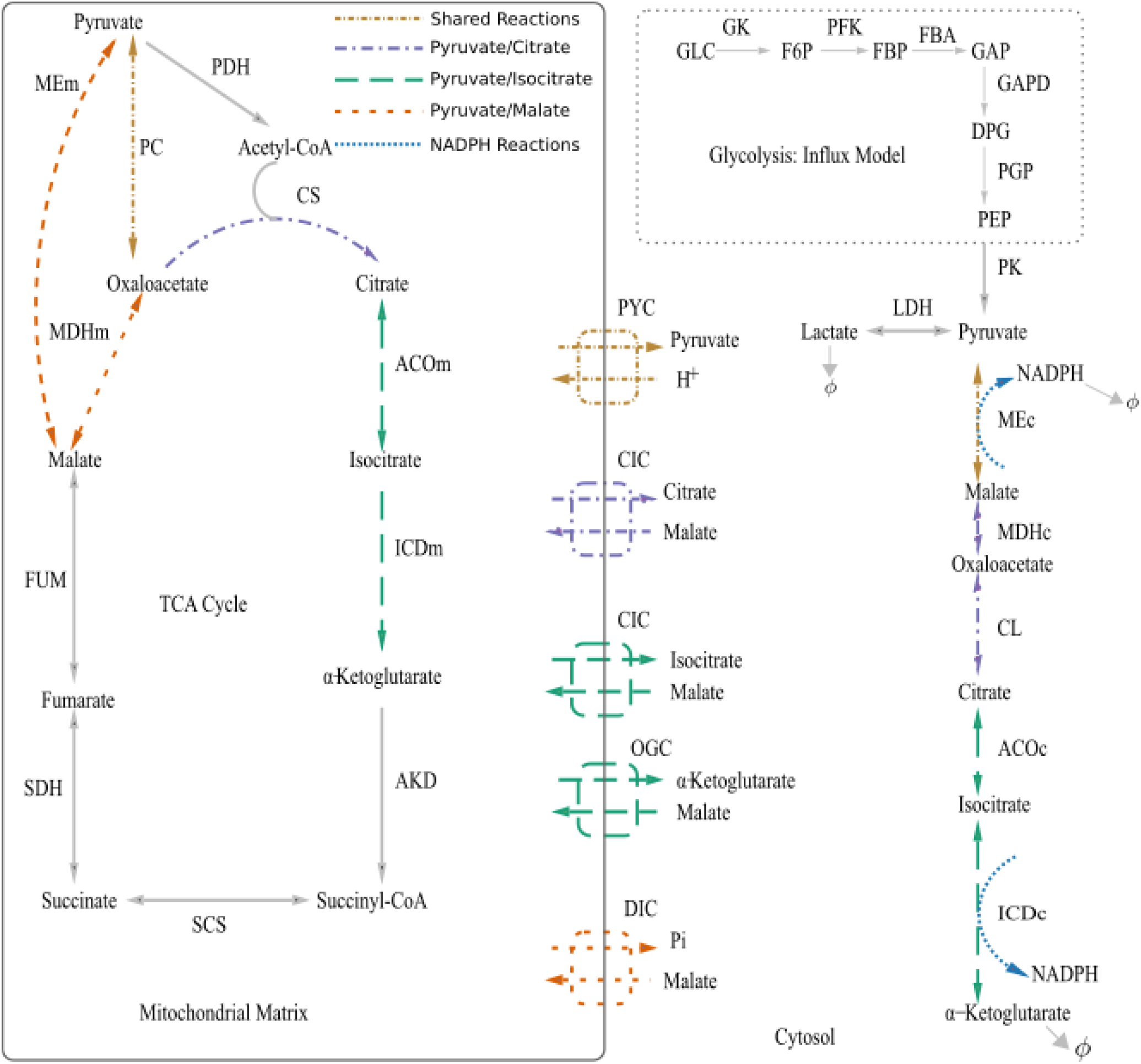
**Pyruvate cycling pathways in β-cells: Shared reactions (brown):** Pyruvate is converted to oxaloacetate by PC. Pyruvate is transported between the cytosol and mitochondria through the pyruvate transporter (PYC). In the cytosol, malate is converted to pyruvate by malic enzyme (MEc). **Pyruvate/malate shuttle (orange)**: Oxaloacetate is converted to malate, while malate is converted to pyruvate through mitochondrial malic enzyme or exported to the cytosol through the dicarboxylate carrier (DIC), where cytosolic malic enzyme converts malate to pyruvate. **Pyruvate/citrate shuttle (violet)**: oxaloacetate is converted to citrate, then transported into the cytosol, where citrate lyase converts citrate to oxaloacetate, which is then converted to malate by cytosolic malate dehydrogenase (MDH_c_). Finally, pyruvate is formed. **Pyruvate/isocitrate shuttle (green)**: Oxaloacetate is converted to citrate and then to isocitrate, both of which are transported into the cytosol. Citrate is converted to isocitrate by ACO_c_. Isocitrate can be converted to α-ketoglutarate (α-KG) via ICD_c_. Isocitrate is then converted to α-KG by ICD_c_. Then α-KG can enter the mitochondria for conversion to malate by TCA cycle enzymes, and subsequent conversion to pyruvate by ME_m_ or ME_c_, thus completing the pyruvate cycle. **NADPH** producing reaction steps are labeled in red. The six-step influx glycolysis model (boxed) is excluded from the analysis.

PC-based pyruvate cycling involves regeneration of pyruvate from TCA cycle intermediates via three distinct pathways (Figure 1): the pyruvate/malate cycle, the pyruvate/citrate cycle, and the pyruvate/isocitrate cycle. Each cycle begins with the conversion of mitochondrial pyruvate (PYR_m_) to oxaloactetate (OAA_m_) and ends with the conversion of malate to pyruvate by malic enzyme. (Malic enzyme is active in both the mitochondria and the cytosol. In β-cells, the cytosolic form carries most of the pyruvate cycling flux [17]).

Recent studies have focused on the identification of the metabolic coupling factors (MCF) that act as signals in the amplifying pathway. These studies provide growing evidence that the pyruvate-cycling pathways generate a metabolic factor that couples increased glucose consumption to insulin release [8], [18]–[20]. A number of MCFs have been proposed, including NADPH, α-ketoglutarate (or its derivatives), and guanosine-5’-triphosphate (GTP) (generated by succinyl-CoA dehydrogenase (SCS)) [1], [21]. Recent observations suggest that NADPH is a key signalling molecule. NADPH is a by-product of all of the pyruvate cycling pathways [22]; it is generated by malic enzyme (a step shared by all three cycles), and by isocitrate dehydrogenase, which is active in the pyruvate/isocitrate pathway. Isocitrate dehydrogenase silencing was found to inhibit GSIS [8].

Several groups have produced computational models describing aspects of the TCA cycle and GSIS. Westermark and colleagues [5] developed a model of mitochondrial nicotinamide adenine dinucleotide (NADH) shuttling (involving 10 metabolites and 19 enzymatic reactions). They validated the model against the findings of Eto *et al*. [23], which characterize the NADH shuttle in β-cells. The TCA cycle has been the subject of many modelling studies. A detailed model of mitochondrial metabolism was developed by Yugi and Tomita [6]. Their model describes 58 enzymatic reactions involving 117 metabolites, and incorporates four pathways: the respiratory chain, the TCA cycle, fatty acid β oxidation, and the inner membrane transport system. Jiang *et al*. [24] developed a detailed model of GSIS that describes 44 enzymatic reactions and 59 metabolic state variables. Their model describes five metabolic pathways: glycolysis, the TCA cycle, the respiratory chain, NADH shuttling, and the pyruvate cycle. While these studies involve validation against a range of experimental findings, a systematic corroboration with experimental results on pyruvate cycling, as presented here, has not been attempted. Furthermore, we perform a global sensitivity analysis on the pyruvate cycling pathways to identify control points that could not have been identified by local sensitivity or similar linear analysis methods.

### Model Description

We extended the previous modelling efforts by developing a mathematical model of β-cell metabolism that describes pyruvate cycling. Our model describes the TCA cycle, the pyruvate/malate shuttle, the pyruvate/citrate shuttle, and the pyruvate/isocitrate shuttle, as shown in Figure 1. The model describes 24 metabolites involved in 30 enzymatic reactions; it draws elements from the models of Yugi and Tomita [6], and Westermark *et al*. [5]. Next, we used kinetic mechanisms available from the SABIO-RK database [25]–[26] to model glycolysis influx. The model involves 129 parameters. Ninety-five of the parameter values were obtained directly from literature. The remaining 34 were calibrated by fitting to the experimental observations of Ronnebaum *et al*. [8]. We tested the model’s accuracy by comparing model predictions to qualitative and quantitative observations of system behaviour as reported in the literature on β-cell metabolism [8]–[16]. We then carried out local and global sensitivity analysis to identify important control points in the pyruvate cycling pathways.

The ordinary differential equation model describes the kinetics of 30 enzyme-catalyzed reactions in β-cell metabolism (Figure 1). The model’s state variables are the dynamically independent concentrations of 24 metabolite species in two compartments—the mitochondrial matrix and the cytosol. The mitochondrial inter-membrane space is neglected. One additional species concentration (NADPH) is determined through conservation. Details of the model kinetics and parameter values are described in the supplementary text.

The network input is extra-cellular glucose, which enters the cytosol via the high-capacity, low-affinity glucose transporter-2 (GLUT2). This transport step is modelled as previously reported by Sweet and Matschinsky [27]. Glucose is converted to pyruvate by the six-step glycolysis pathway. The kinetics of all glycolytic reactions were drawn from the SABIO-RK database [26], with some adjustments (details in the supplementary text). The glycolytic pathway is treated as a fixed influx module and is not included in the model analysis presented below. The end-product of glycolysis, cytosolic pyruvate, is either converted to lactate by the lactate dehydrogenase (LDH) or transported into the mitochondrial matrix. The former process carries much less flux than the latter; LDH activity is weak in β-cells [9]. The TCA cycle operates within the mitochondrial matrix. All components of the three pyruvate cycling processes—the pyruvate/malate cycle, the pyruvate/citrate cycle, and the pyruvate/isocitrate cycle—are included in the model.

The **pyruvate/malate cycle** involves conversion of mitochondrial oxaloactetate OAA_m_ to malate, via mitochondrial malate dehydrogenase (MDH_m_). Mitochondrial malate then follows one of two routes: it can be directly converted to pyruvate by mitochondrial malic enzyme (ME_m_), or it can be transported to the cytosol via the dicarboxylate carrier (DIC) and then converted back to pyruvate by cytosolic malic enzyme (ME_c_).

The **pyruvate/citrate cycle** begins with OAA_m_ following its normal route through the TCA cycle: it combines with acetyl-CoA to form citrate via citrate synthase (CS). This mitochondrial citrate can then be converted to isocitrate by mitochondrial aconitase (ACO_m_). Mitochondrial citrate and isocitrate are transported into the cytosol by the citrate/isocitrate carrier (CIC). The cytosolic form of aconitase (ACO_c_) then converts isocitrate to citrate. Citrate lyase (CL_c_) converts citrate to oxaloacetate releasing acetyl-CoA. This cytosolic OAA_c_ can then be converted to malate by the cytosolic form of malate dehydrogenase (MDH_c_). Finally, malate is converted to pyruvate by malic enzyme, thus completing the cycle. This last step is shared with the pyruvate/malate cycle.

The **pyruvate/isocitrate cycle**, like the pyruvate/citrate cycle, starts with oxaloacetate being converted to citrate and isocitrate and the subsequent exit of these metabolites from the mitochondria through the citrate/isocitrate carrier (CIC). In the cytosol, citrate is converted to isocitrate by ACO_c_. Isocitrate is then converted to α-ketoglutarate (α-KG) by cytosolic NADP^+^-dependent-isocitrate dehydrogenase (ICD_c_). α-KG is then transported back into the mitochondria by the oxoglutarate carrier (OGC). Once in the mitochondria, α-KG follows the TCA cycle to be converted to malate, which can then be converted back to pyruvate by malic enzyme. This last step is shared with the other cycles.

#### Simulation method

Simulations were carried out in MATLAB® (function ode15s). To ensure the simulations reached steady state, we used a trust-region root finding method to confirm the steady state conditions (MATLAB^®^ function fsolve). Details of simulations are provided in the supplementary text.

#### Parameterization Approach

The reaction kinetics, as well as the bulk of the parameter values, were derived from the previous models of Westermark *et al*. [5], and Yugi and Tomita [6] and the SABIO-RK [25], [26], and Brenda databases [28], [29]. After formulating the model and identifying nominal parameter values, we performed a preliminary sensitivity analysis from which we identified a set of 34 parameters that were consistently ranked highly by all sensitivity ranking methods. We fixed the other 95 parameters at values obtained from the literature (details in the supplementary text), and then fit the values of the 34 high-sensitivity parameters (the resulting values remained near the nominal values obtained from the literature, details in the supplementary text). This calibration was performed by fitting to the experimental observations of Ronnebaum *et al*. [8]. That paper provides 32 steady state metabolite concentration measurements (eight species under four experimental conditions). We will refer to this data as the training set. Parameter values were estimated using the simplexSB and simannealSB algorithms provided by the SBTOOLBOX2 software package [30], which minimize the sum of squared errors, weighted by the variability in the data. Details of the fitting procedure, including the bounds used for the parameter search, are reported in the supplementary text. Having found a best-fit to the training set, we then verified the model by comparing its predictions against a range of experimental results on β-cell metabolism, as described in the next section. None of this test set data was used for fitting. We have made the source codes to reproduce model results available at https://github.com/r2rahul/pyruvatecycle following the guidelines for publishing reproducible models described by Tiwari *et al*. [31]–[33]. Further details are provided in the supplementary text.

### Model Analysis Methods

A primary aim of this study is to identify network *control points* that have significant influence over the rate of pyruvate cycling and consequent NADPH production. Our goal is to identify parameters whose values have significant influences over the key metabolite concentrations (pyruvate and NADPH), the pyruvate cycling ratio (defined as the ratio of PC flux to the sum of all TCA cycle fluxes, more details provided in supplementary text section 2), and the flux through the pyruvate cycling reactions. These highly significant parameters can serve as points of investigation for further experimentation, with the goal of identifying effective drug targets for treatment of type 2 diabetes.

To identify network control points, we employed local and global sensitivity analysis as described next.

#### Local sensitivity analysis (LSA)

We used local parametric sensitivity analysis to investigate the effect of small perturbations in the values of individual parameters. Normalized steady state sensitivity coefficients were calculated using a first order forward difference formula (with 10% deviation from the nominal parameter value). We define *S*_*i*_ as the normalized steady state sensitivity coefficient with respect to parameter *p*_*i*_

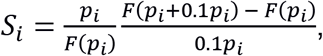

in which *F* is the flux or metabolite concentration of interest. We used Richardson extrapolation as described in Elden *et al*. [34] to avoid any inaccuracy due to first order forward difference approximation.

#### Global sensitivity analysis (GSA)

Global sensitivity analysis serves two purposes. First, it allows us to explore the nonlinear dependence on model parameters by considering the effects of coincident variations in multiple parameters. Second, it allows a wide exploration of parameter space, which is especially important given the uncertainty of the estimates of the nominal parameter values. Following, the recommendation of Zhang and Rundell [35], we used two complementary global sensitivity analysis methods: Partial rank correlation coefficient (PRCC) analysis (with Latin hyper-cube sampling) [36] and the extended Fourier amplitude sensitivity test (eFAST), a variance-based method [37]–[39]. Both methods are implemented in SBTOOLBOX2 [30], [40].

Variance-based global sensitivity analysis methods address both single-parameter variation—called first-order sensitivity—and multi-parameter variation—called total-effect sensitivity. Comparisons between the first-order and total-effect sensitivities provide insight into nonlinear interactions among parameter-dependent effects. In addition to the sensitivity of key model outputs (fluxes and steady state concentrations) we also considered the sensitivity of an overall system output, defined as the sum of the squared deviation (from nominal) in all steady state metabolite concentrations. A complete description of the objective function is provided in the supplementary text.

## Results

### Model training

The model was calibrated against a training set provided by the experiments in Ronnebaum *et al*. [8], in which siRNA specific for the mRNA of ICD_c_ was used to reduce the activity of ICD_c_. To simulate the effect of the siRNA treatment, we reduced the V_max_ parameter of ICD_c_ by the measured decrease (by 39.1%) in enzyme activity. Ronnebaum *et al*. [8] collected steady-state measurements of the concentrations of eight metabolites in each of four cases: control and knock-down at low glucose (3mM) and high glucose (12mM). Simulating these four cases to steady state, we fit the model by minimizing a sum of squared errors measure of the difference between the steady-state model predictions and the observed values. Details of the computation are included in the supplementary text. The best-fit model behavior is shown in Figure 2. The best-fit model parameters are reported in the supplementary text, along with the values of the remaining model parameters, which were taken from the models of Yugi and Tomita [6] and Westermark *et al*. [5], and the Brenda [28], [29] and SABIO-RK [25], [26] databases.

**Figure 2:**
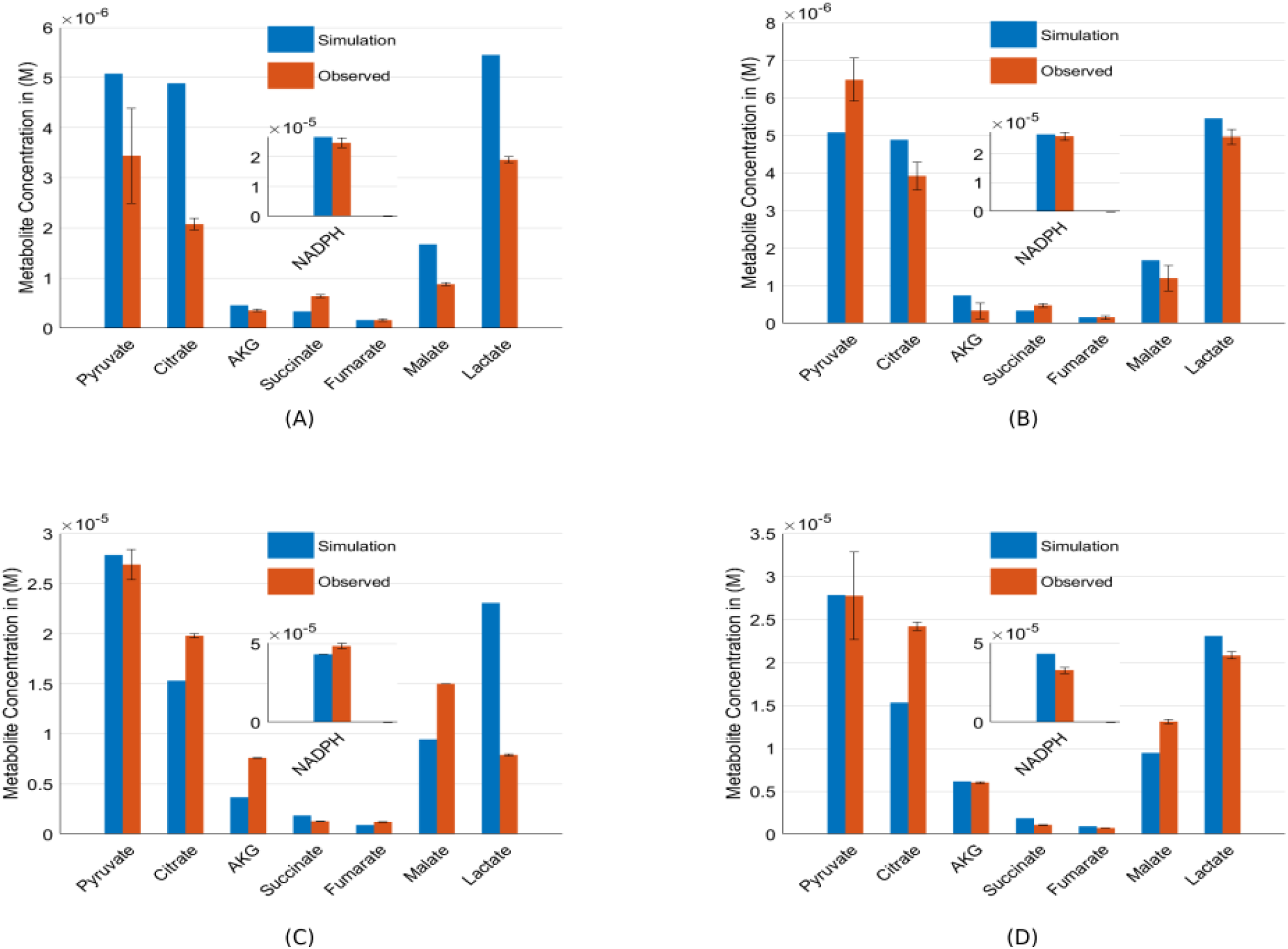
Best-fit model steady-state model predictions compared to data from Ronnebaum *et al*. [8]. **A**. Low glucose, control. **B**. High glucose, control. **C**. Low glucose, ICD_c_ knock-down by 39.1%. **D**. High glucose, ICD_c_ knock-down by 39.1%. NADPH inset is on a different scale.

#### Model Testing

We validated the model by comparing model simulations to a series of observations gathered from the literature, as described below. None of these datasets were used in parameter fitting.

### Lactate dehydrogenase activity

The model corroborates the experimental observation of Sekine *et al*. [9] that, in comparison with other cell types, β-cells exhibit significantly reduced lactate dehydrogenase (LDH) activity. In the model network, glucose-derived pyruvate is either converted to lactate by LDH or is transported into the mitochondria by the pyruvate transporter (PYC). We compared the flux through these reactions over a wide range of glucose levels (2.5 mM to 22 mM) and found that LDH exhibits between 2-4% of the PYC flux.

### Malic enzyme activity

It has been shown that in β-cells, the cytosolic form of malic enzyme (ME_c_) contributes approximately 90% of the total malic enzyme activity in the cell, at glucose concentrations ranging from 3 mM to 20 mM [10]. Simulations of the model predict that roughly 95% of malic enzyme flux is carried by ME_c_, for glucose levels ranging between 2.5 mM and 22 mM.

### Pyruvate carboxylase activity

Radio-isotopic experiments have revealed that in β-cells approximately 40% of glucose-derived pyruvate enters the TCA cycle via PC-catalyzed conversion to OAA, with the remainder metabolized to acetyl-CoA via pyruvate dehydrogenase [1]. The model predicts that over the range of 12-20 mM glucose, approximately 40% of pyruvate enters the TCA cycle through pyruvate carboxylase. This percentage increases to above 50% at low glucose (3 mM).

### Pyruvate cycling

Using a ^13^C NMR isotopomer method, Lu *et al*. [16] measured the rate of pyruvate cycling, which they defined as the ratio of PC flux to the overall TCA cycle flux. They found that the anapleurotic flux catalyzed by PC is correlated with GSIS, while pyruvate dehydrogenase mediated entry of pyruvate into the TCA cycle is not significantly affected by changes in glucose abundance. We compared the model predictions of the percentage increase in the pyruvate cycling ratio at 3, 6, and 12 mM glucose as observed in the experiments of Lu *et al*.. When glucose is increased from 3 to 6 mM, the observed percentage increase in pyruvate cycling ratio was 117% while model simulation shows a 56% increase. When the glucose concentration was raised from 6 to 12 mM, the observed increase in the ratio was 65% while simulations predict an increase of 49%. While the model underestimates the cycling ratio, it correctly captures the increase in cycling ratio with glucose availability. The experiments of Lu *et al*. also revealed that the acetyl-CoA concentration does not increase linearly with glucose availability. When glucose concentration is raised from 3 to 6 mM, the model predicted a 29% increase in acetyl-CoA concentration, whereas a 41% percent increase was observed. Similarly, the model predicted a 16% increase in acetyl-CoA concentration when glucose concentration is raised from 6 to 12 mM, while the observed increase was 70%. Finally, an increase in the glucose concentration from 3 to 12mM was observed to cause an 86% increase, while the model predicted only a 40% increase. Overall, the model underestimates the percentage increases, but correctly captures the acetyl-CoA saturation trend.

Ronnebaum *et al*. [8] studied the effect of an ICD_c_ knock-down on pyruvate cycling; they measured the pyruvate cycling ratio in wild type and ICD_c_ knock-down strains. They observed that knockdown of ICD_c_ RNA by 39% caused a reduction of pyruvate cycling to 84±4% of the wild-type ratio at high glucose (12 mM), while at low glucose (3 mM), the knock-down resulted in a slight increase in pyruvate cycling ratio. Our simulations of the knockdown (reducing the V_max_ of ICD_c_ by 39%) showed no significant effect on the pyruvate cycling ratio at low or high glucose levels.

### Citrate lyase knock-down

Joseph *et al*. [12] conducted siRNA-mediated suppression of citrate lyase (CL), reducing activity by 75±4%. This resulted in a 52±7% reduction in cytosolic oxaloacetate, and no significant impact on the NADPH:NADP^+^ ratio (in steady state, at 16.7 mM glucose). We simulated this experiment by reducing the V_max_ value of CL by 75%. At steady state, at 16.7 mM glucose, the resulting change in the NADPH:NADP^+^: ratio is .01%; the cytosolic oxaloacetate concentration drops by 62.8%. The model predicts a more modest drop in OAA_c_ at lower glucose levels, e.g. 54.4% at 3 mM glucose.

### Pyruvate carboxylase knock-down

Jensen *et al*. [13] conducted an siRNA-mediated knock-down of pyruvate carboxylase. In their knock-down strain, PC activity was reduced by 65%, which resulted in a negligible effect on the NADPH:NADP^+^ ratio. Our model prediction, corresponding to a 65% drop in the V_max_ parameter of PC, shows a 9.9% decrease in the NADPH:NADP^+^ ratio.

Jensen *et al*. also analyzed the effect of the PC knockdown on the concentrations of TCA cycle intermediates. They found that the concentrations of succinate, malate, and citrate were unaffected by the PC knock down. Our model simulation of the knockdown predicts succinate, malate, and citrate are reduced by 0.01%, 5.1%, and 0.11% respectively. Furthermore, Jensen *et al*., found that PC suppression reduced the α-KG concentration by 70% at low glucose (2.5 mM) and 31% at high glucose (12 mM). Our simulation follows a similar trend but showed strong reduction of 80% and 60% respectively. Additionally, Jensen *et al*. reported increases in the lactate and acetyl-CoA concentrations after PC knock down. Our simulation captured this trend but showed only nominal concentration increases of 9.91% and 11.67% in lacate and acetly-CoA respectively. Additional model predictions of the effect of varying PC activity are discussed in the sensitivity analysis section.

### Knock-down of cytosolic malic enzyme

To investigate the role of the pyruvate/malate cycle, Ronnebaum *et al*. [14] conducted an siRNA-mediated knock-down of cytosolic and mitochondrial malic enzyme. They found that knock-down of either enzyme by 75% had no effect on the pyruvate cycling ratio at either 2.5 mM or 12 mM glucose. The corresponding model predictions showed a negligible effect of cytosolic malic enzyme knockdown, but a knockdown of mitochondrial malic enzyme had a modest effect: a decrease of 0.08% at 2.5mM glucose but a 9% increase at 12mM glucose.

### Model Trends of NADPH concentration profile

In addition to the measurements of NADPH concentration used for model training (Figure 2), Ronnebaum *et al*. [8] also made measurements of the NADPH:NADP^+^ ratio at a range of glucose concentrations. The model shows reasonable agreement with these findings, as shown in Table 1. However, their observations of the reduction of the NADPH:NADP^+^ ratio in the ICD_c_ knockdown strain (as compared to wild-type) were not captured by the model. They observed a decrease in this ratio of 42% at 3 mM and 17% at 12 mM, while the model simulation showed only a negligible decrease.

**Table 1:** Percentage increase in NADPH concentration in response to an increase in glucose concentration.

| Glucose increment (mM) | Observed (%) | Model Simulation (%) |
| --- | --- | --- |
| 3 to 4 | 33.3 | 16.4 |
| 4 to 6 | 35.7 | 24.6 |
| 6 to 12 | 50.0 | 26.3 |
| 3 to 12 | 125 | 126 |

Sener *et al*. [41] hypothesized that the NADPH:NADP^+^ ratio can be inferred from the pyruvate:malate ratio or the isocitrate:α-KG ratio. Our model prediction shows that, except for a departure in the ICIT:α-KG ratio at low glucose levels, all these three ratios follow very similar trends, as shown in Figure 4. Finally, Hedeskov *et al*. [10] found that the NADPH:NADP^+^ ratio increases 125% when glucose is increased from 3 to 20 mM. The model predicts an increase of 71%.

**Figure 3:**
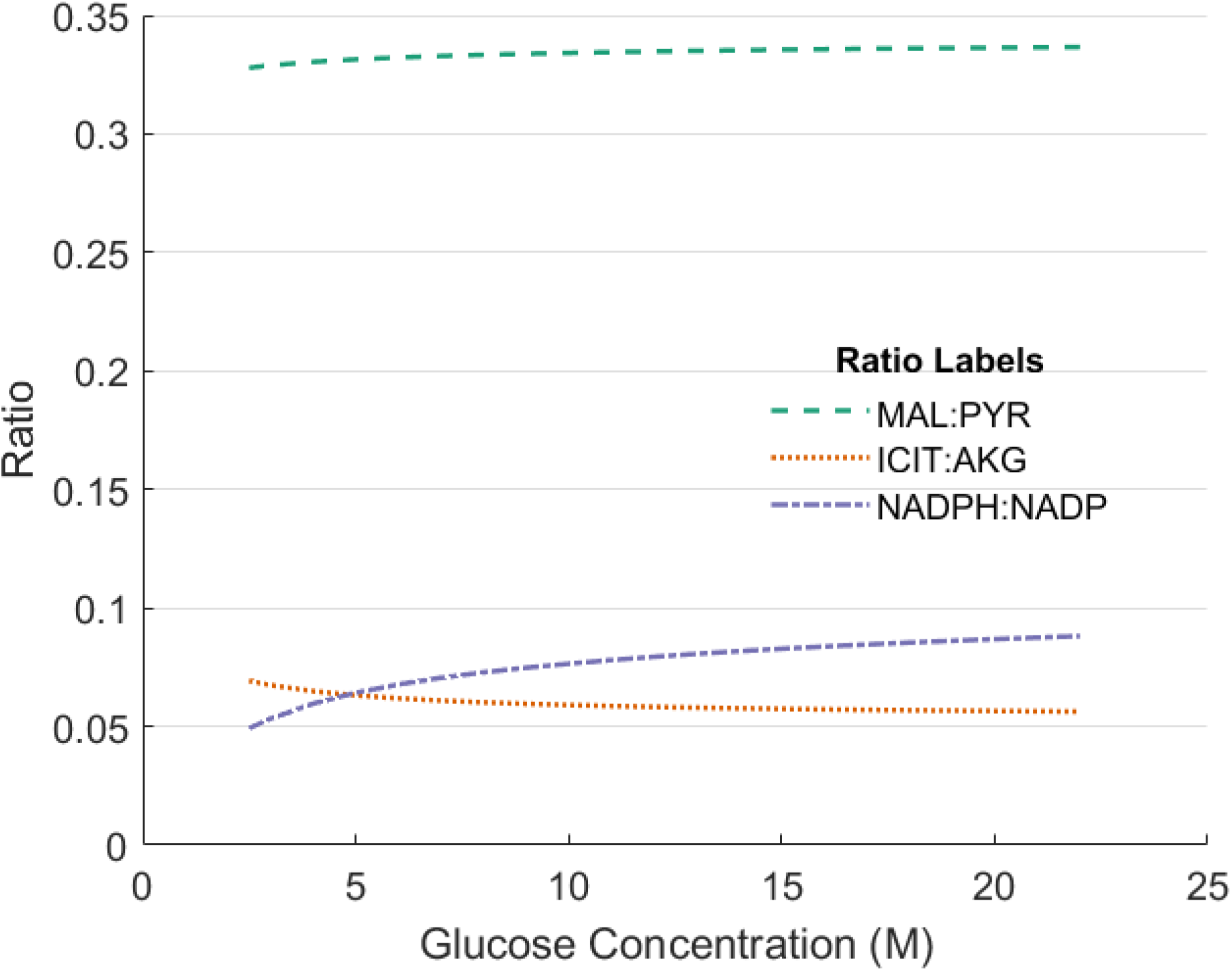
Model simulations of the NADPH:NADP^+^, PYR:MAL, and ICIT:AKG ratios over a range of glucose concentrations.

**Figure 4:**
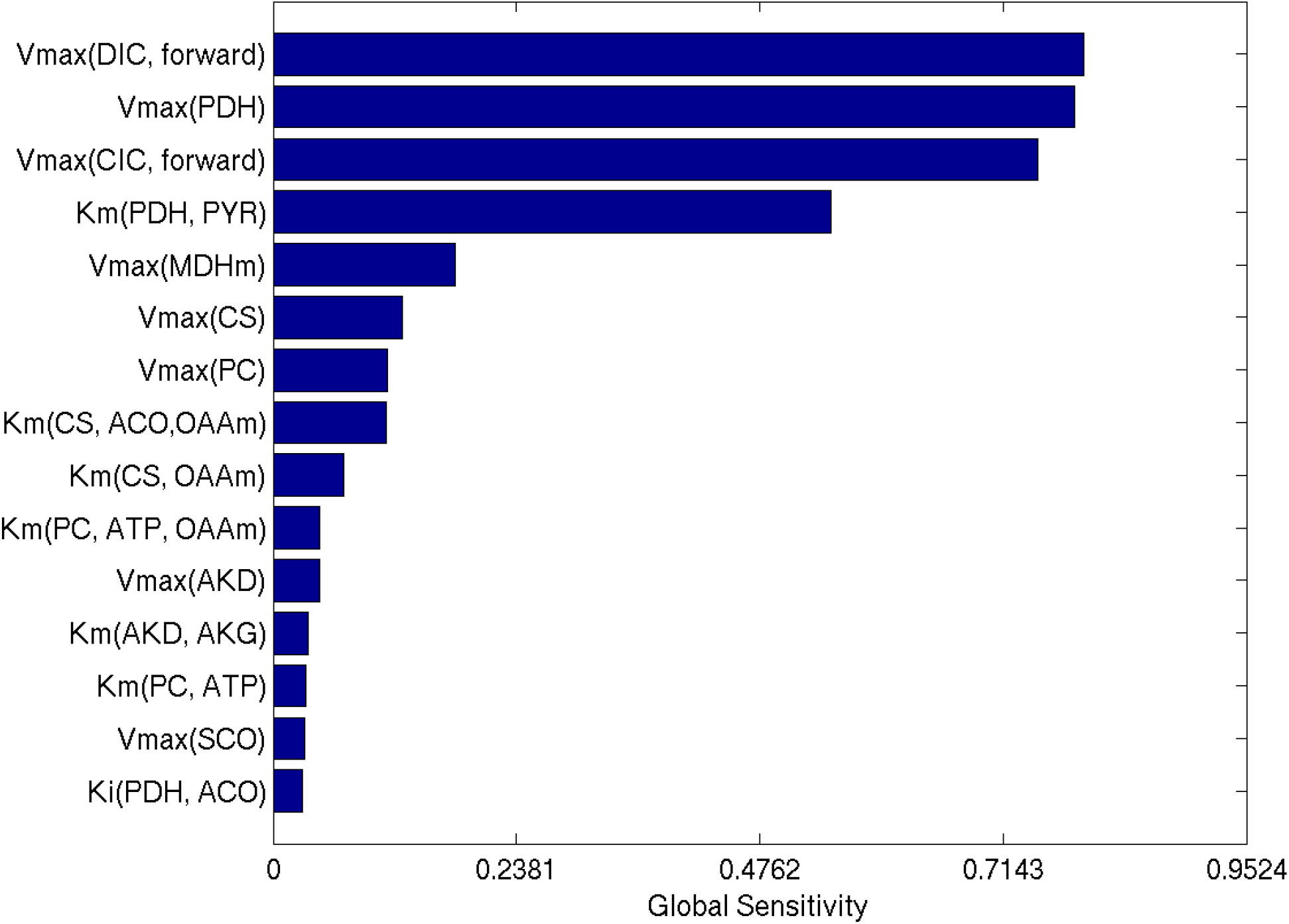
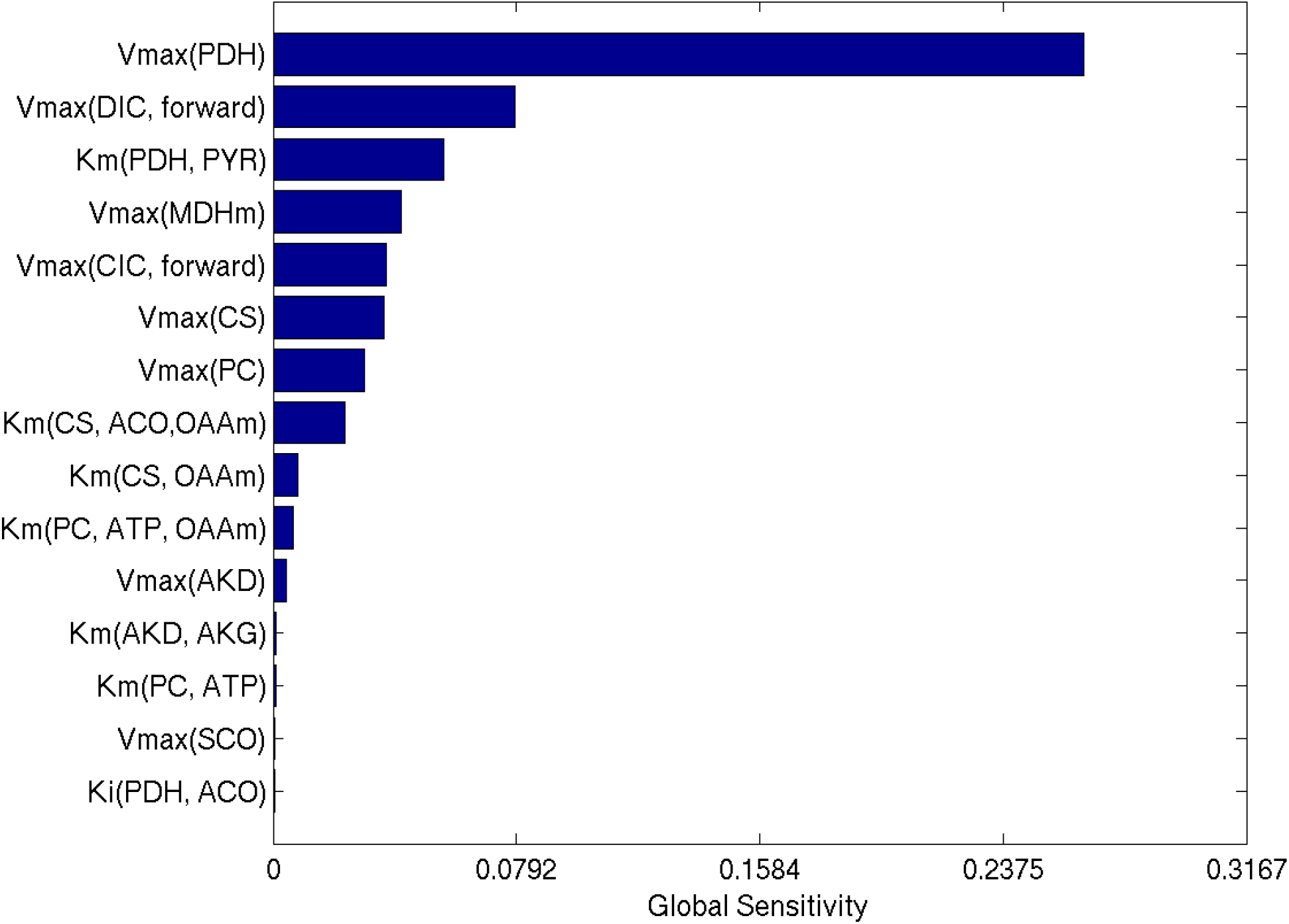
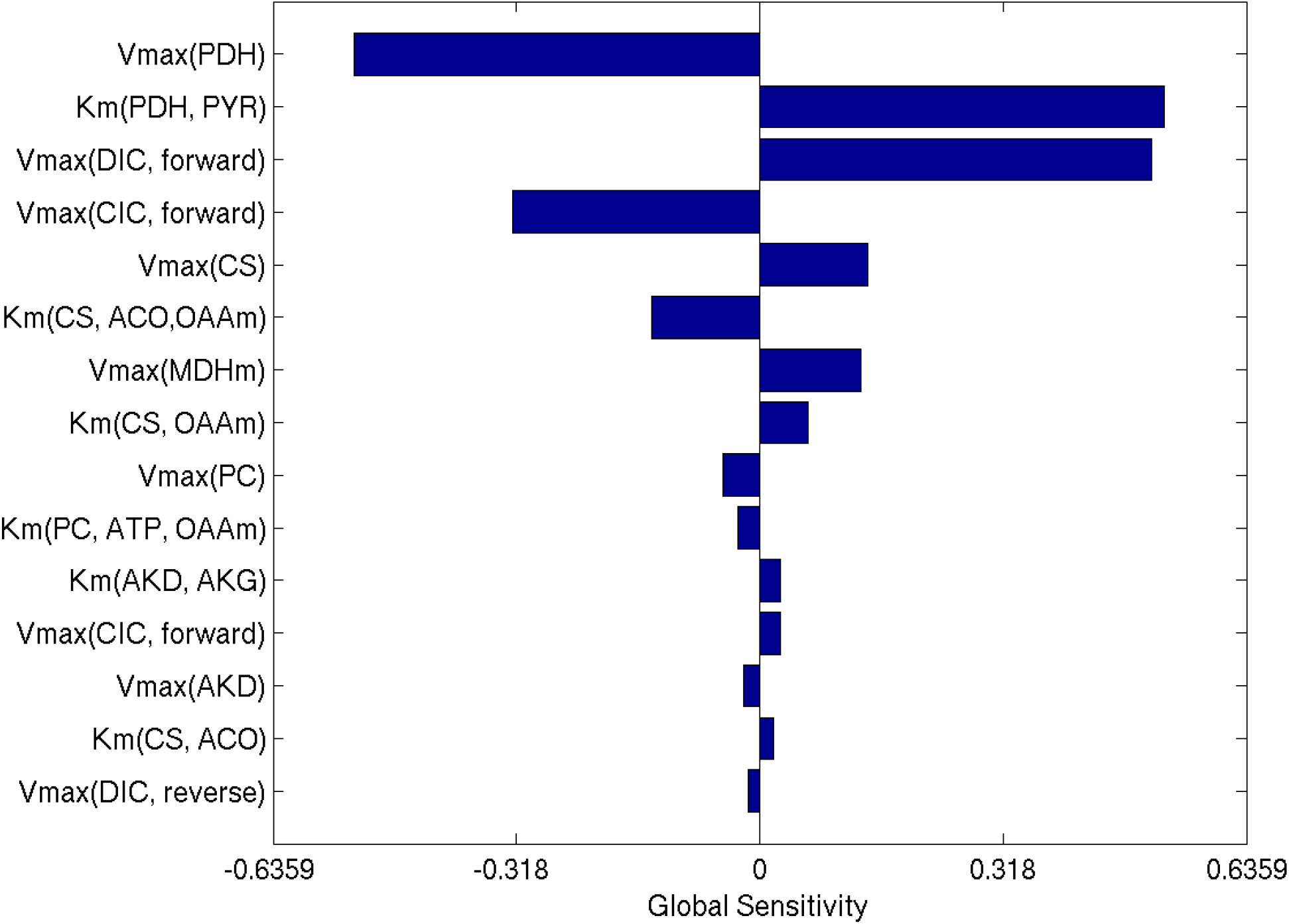
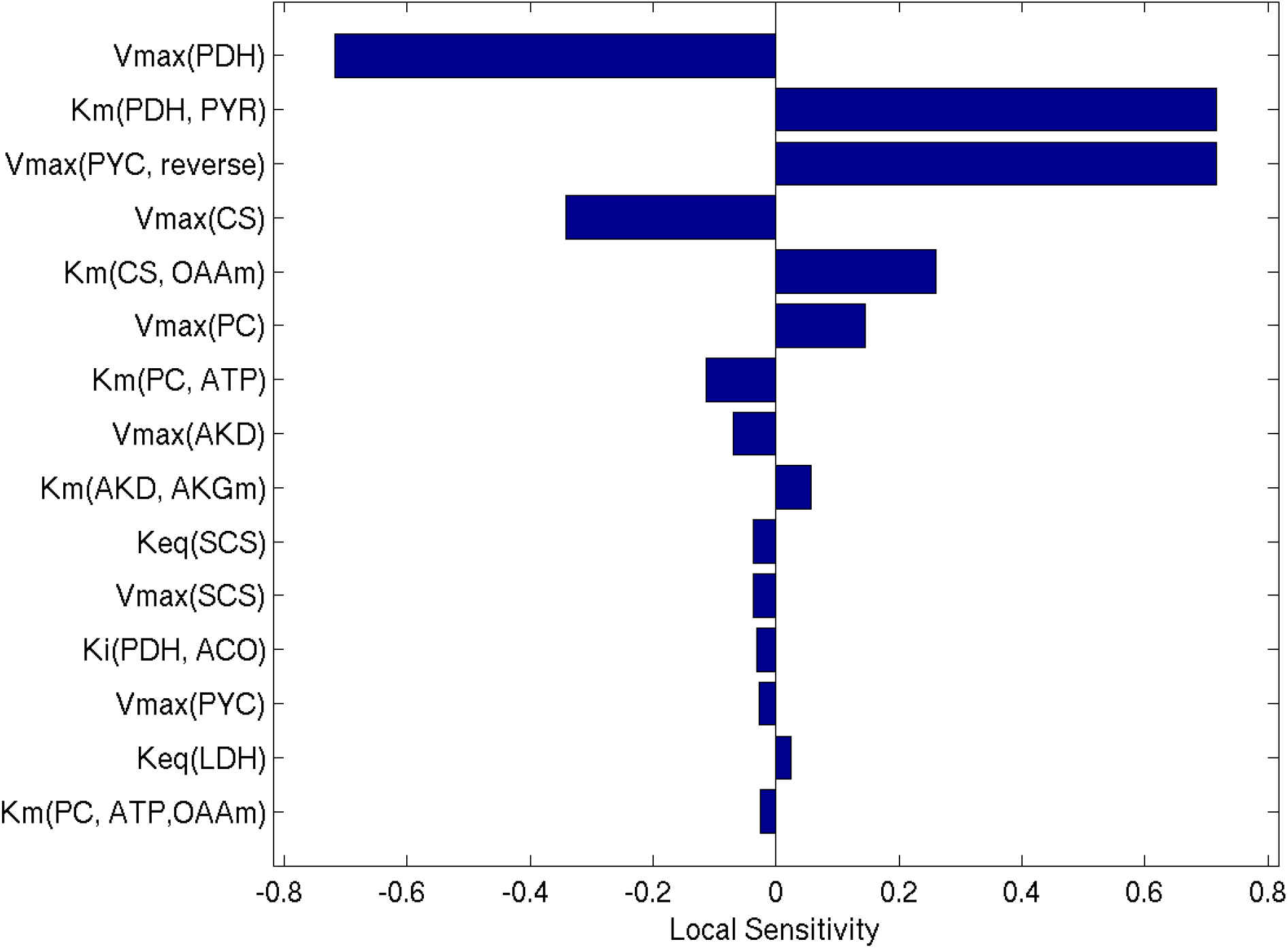
Sensitivity ranking of the pyruvate cycling ratio. **A.** eFAST total effect. **B** eFAST first order. **C.** PRCC. **D.** Local Sensitivity.

## Sensitivity Results

### Sensitivity Rankings

Figures 4 and 5 shows the ranked global sensitivities with respect to the two model outputs that are likely directly related to insulin secretion: the steady-state pyruvate cycling ratio and the steady-state NADPH concentration. Sensitivity rankings of the steady state pyruvate concentration (cytosolic and mitochondrial), the pyruvate carboxylase flux, and the flux through cytosolic isocitrate dehydrogenase are included in the supplementary text.

**Figure 5:**
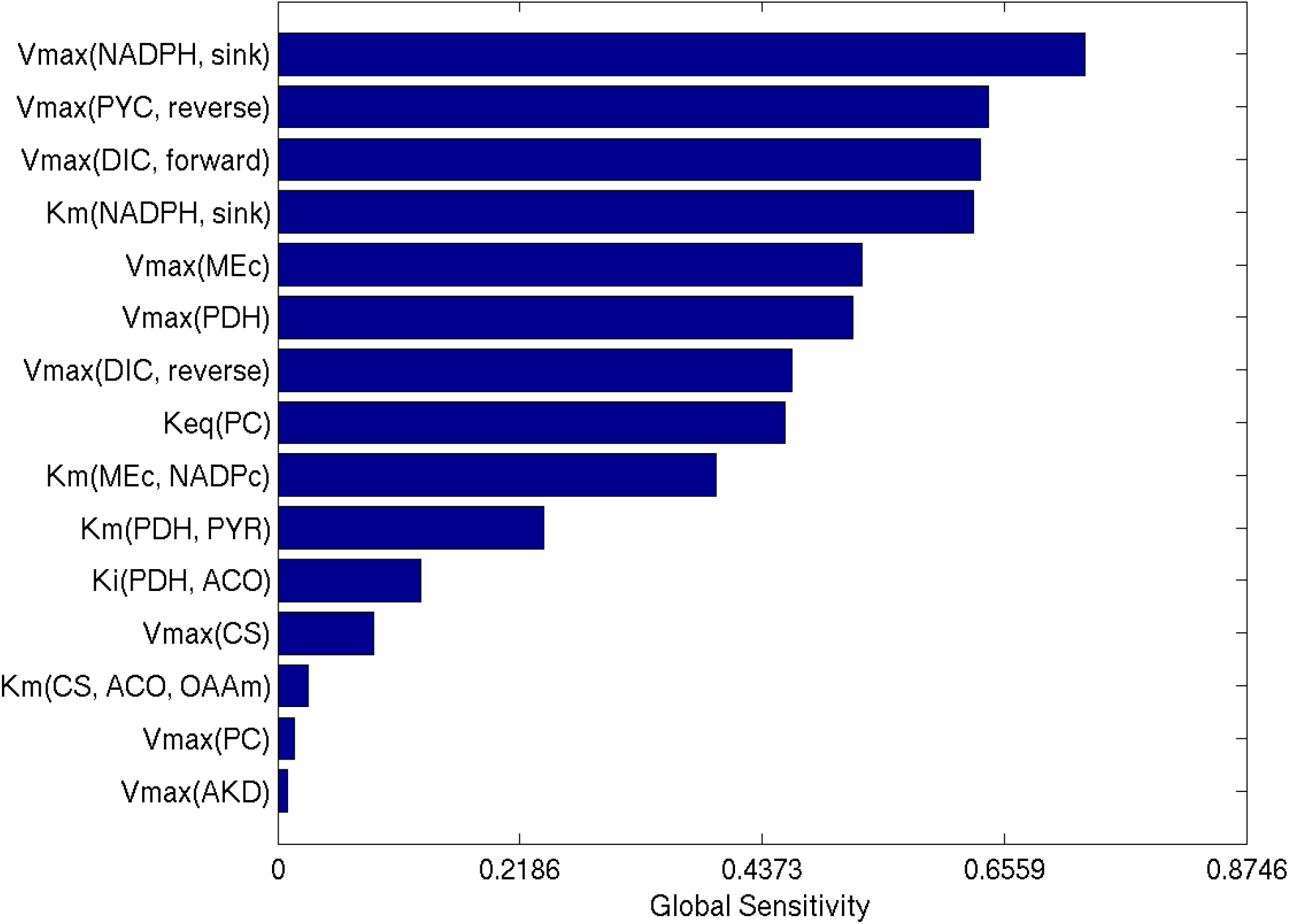
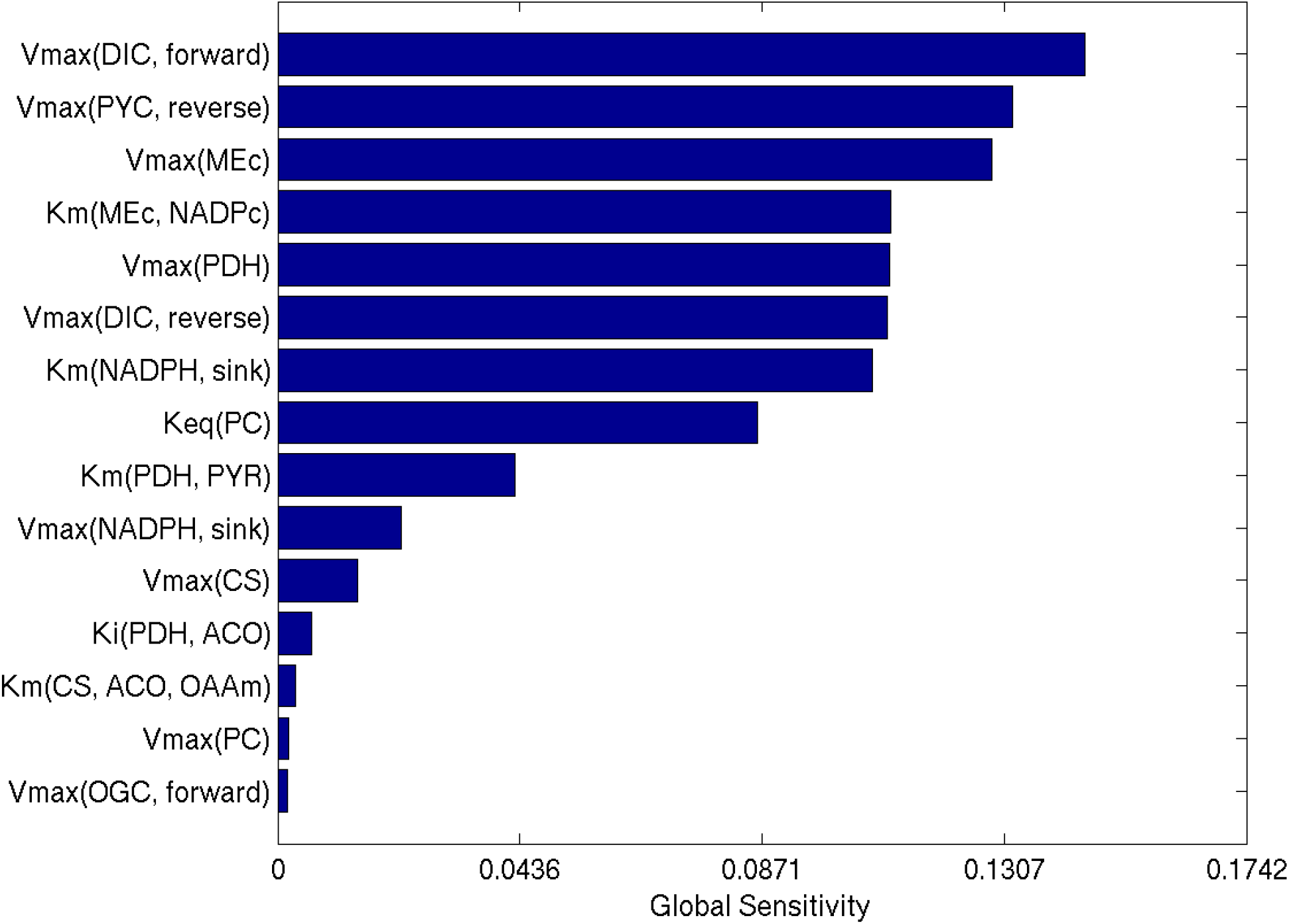
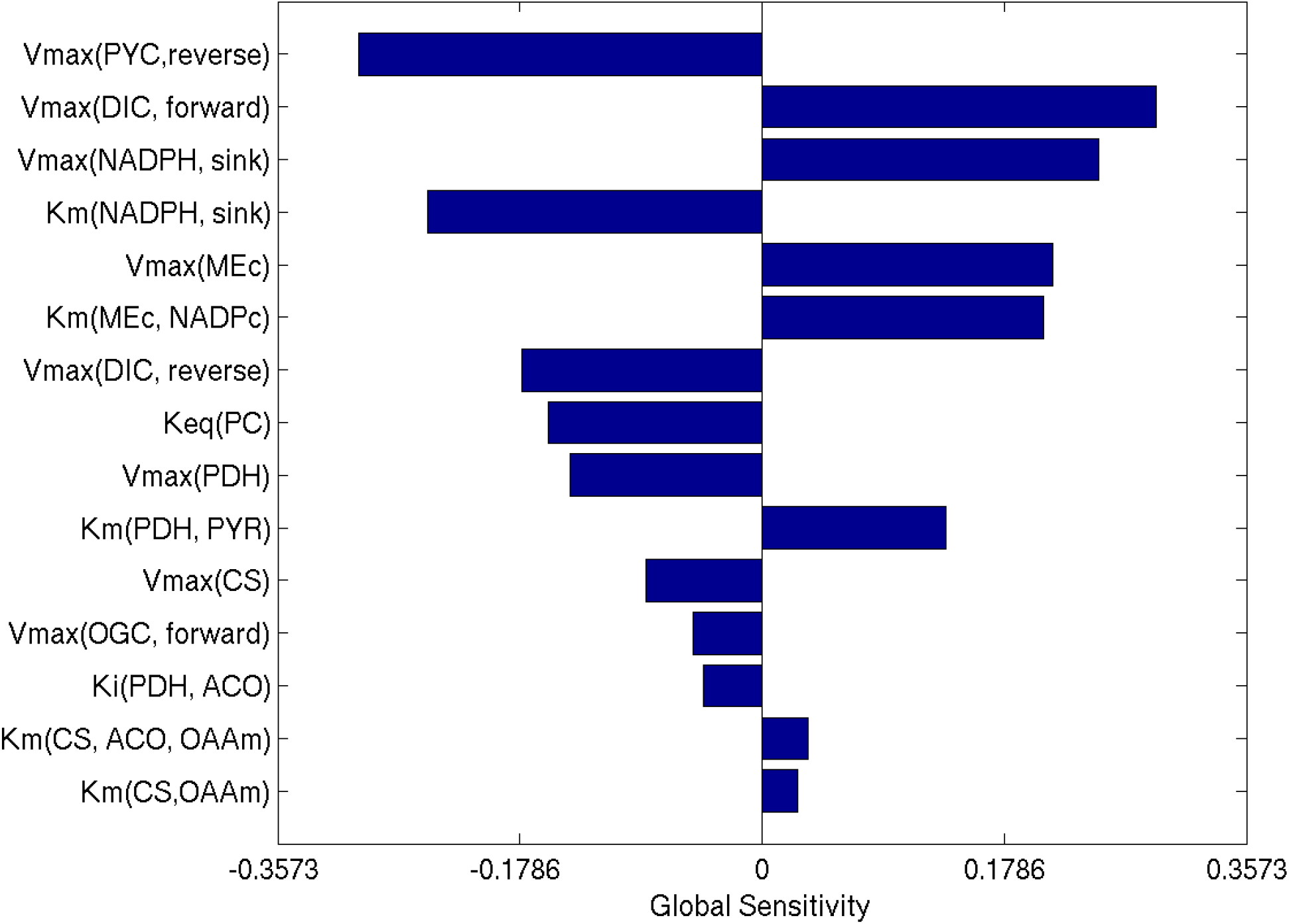
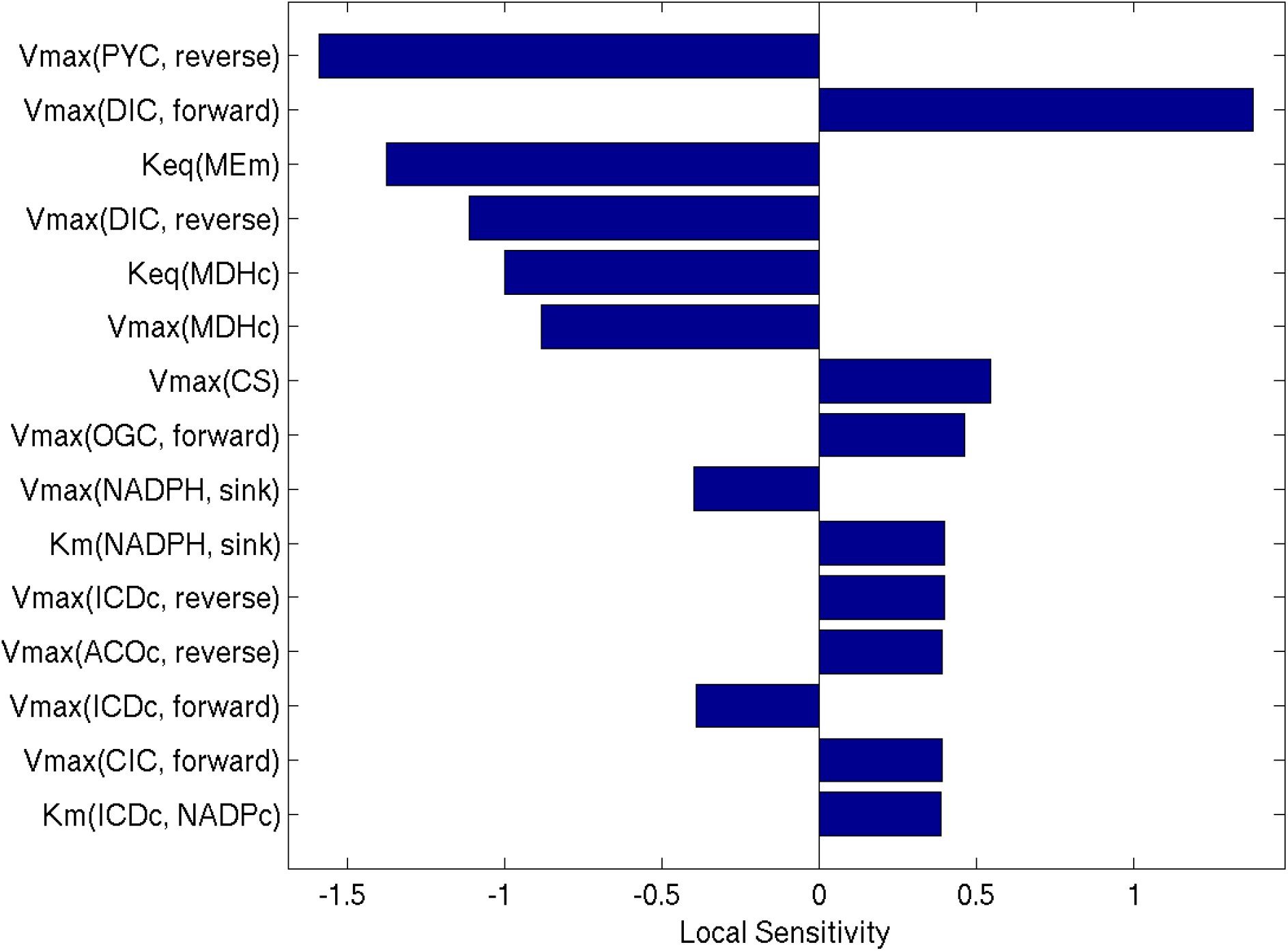
Sensitivity ranking for the NADPH concentration. Panel **A.** eFAST total effect. Panel **B.** eFAST first order. Panel **C.** PRCC. Panel **D.** Local sensitivity.

### Effect of changes in pyruvate carboxylase activity

Because it is the first step in all three pyruvate cycling pathways, pyruvate carboxylase (PC) might be expected to exert significant control over the distribution of flux in the network. We calculated sensitivity measures for the effect of perturbations in the V_max_ of PC on steady state flux for all reactions in the network. Figure 6 shows the five most affected fluxes and the five least affected fluxes.

**Figure 6:**
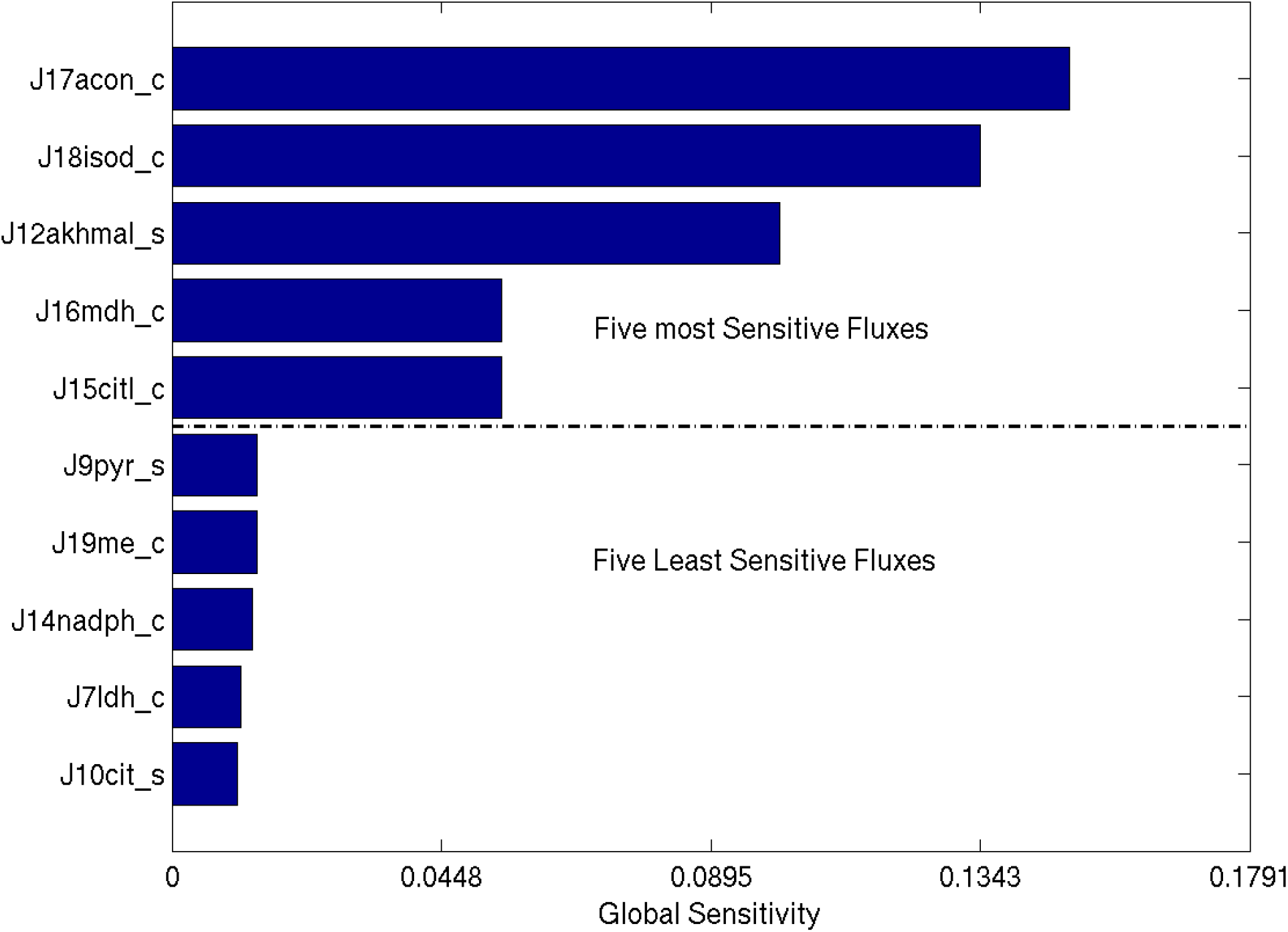
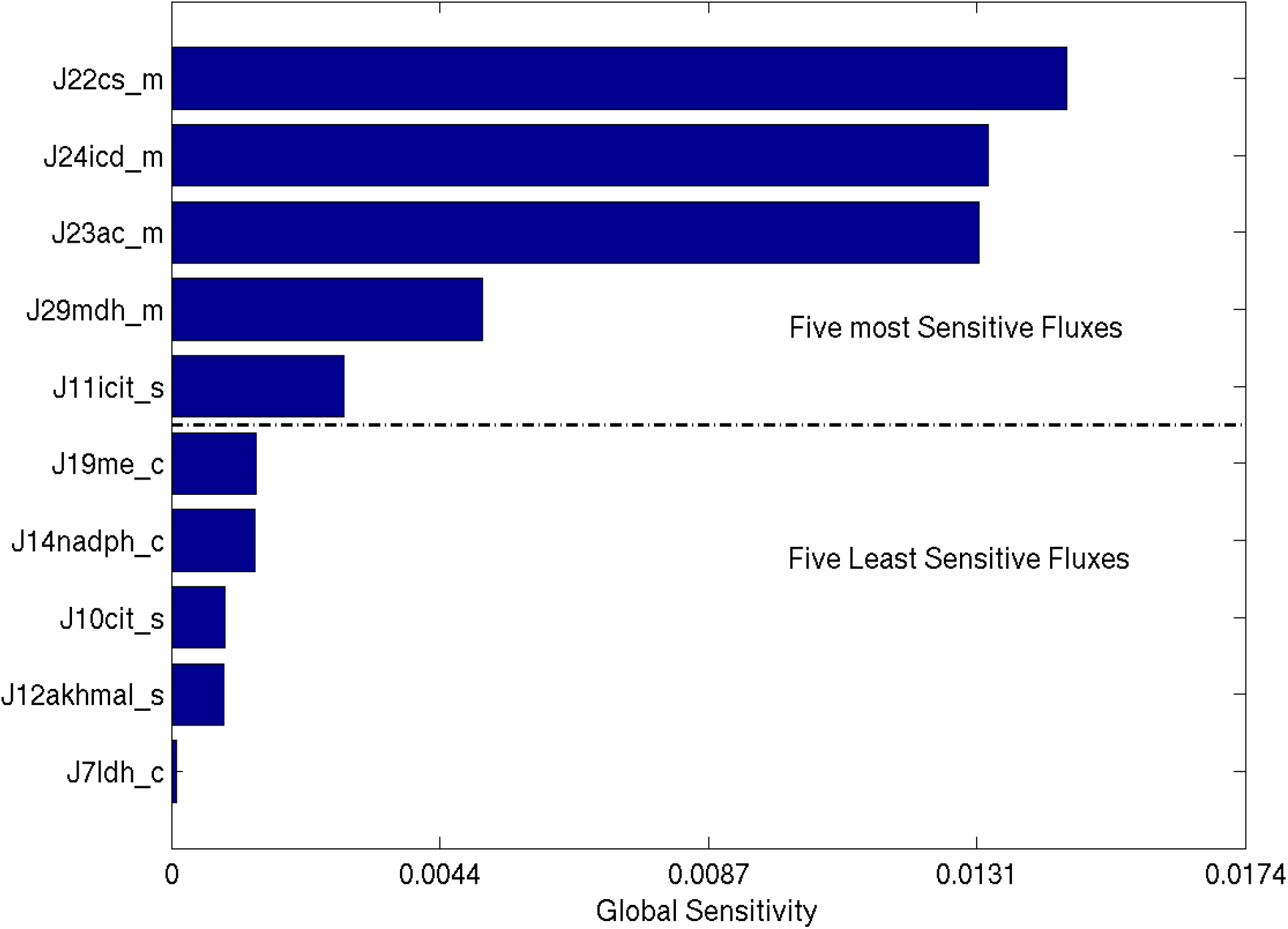
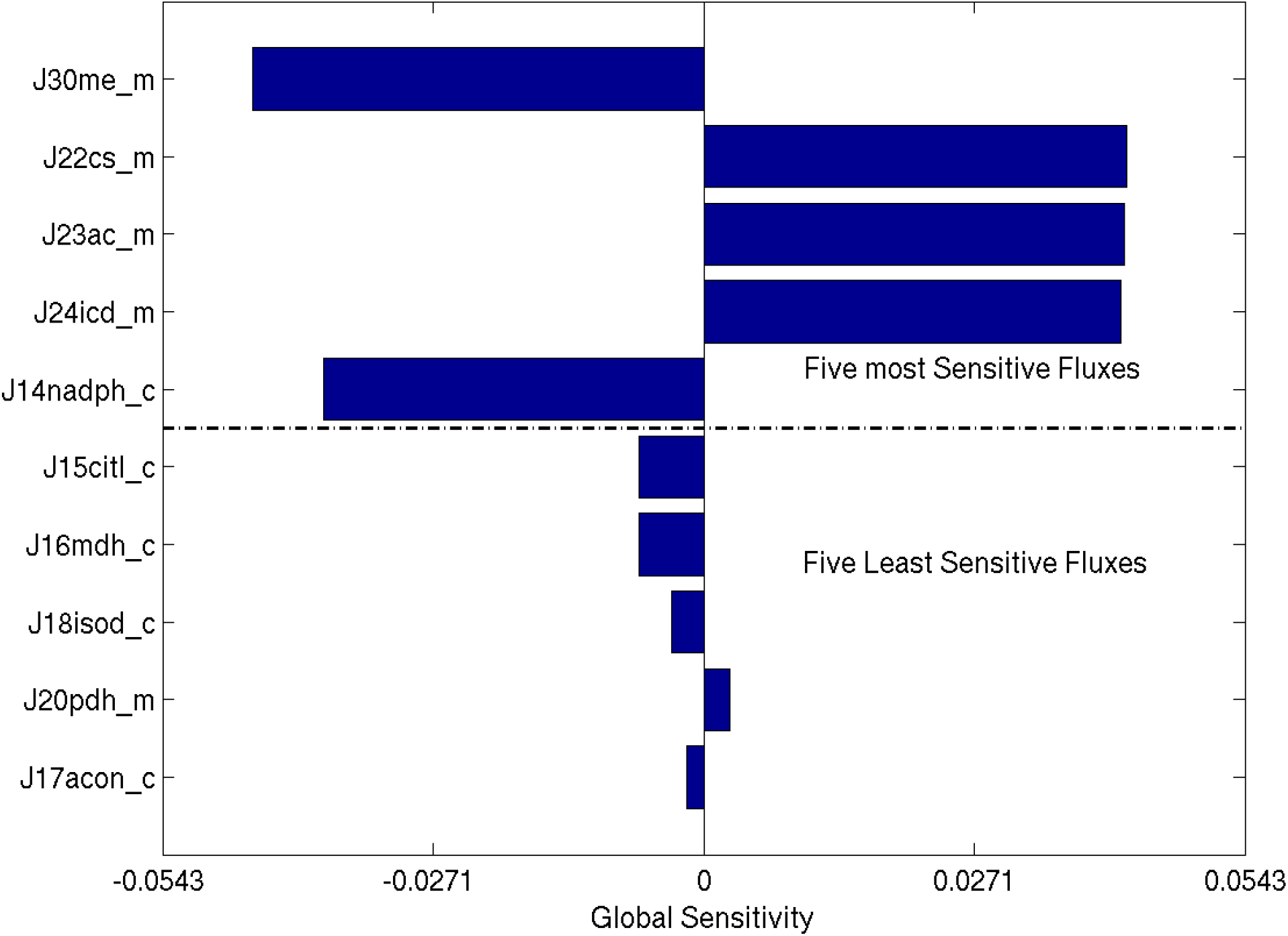
Predicted effect of perturbation in the V_max_ of PC on reaction fluxes. The five most sensitive fluxes and the five least sensitive fluxes are plotted. **A**. local sensitivity. **B**. PRCC ranking, **C.** eFAST total effect and **D.** eFAST first order. For flux descriptions, refer to the supplementary text.

Further we compared the sensitivity rankings of eFAST total-effect at low glucose (3 mM) and high glucose (12 mM). We found that parameters of many pyruvate cycling enzymes, such as V_max_ of ME_c_, are ranked higher at the high glucose concentration compared to their low glucose rankings. See supplementary text for details. These shifts suggest an increase in the activity of pyruvate cycling enzymes at the higher stimulatory glucose concentration.

## DISCUSSION

Given the highly non-linear nature of the model, it is worthwhile to compare the results of different sensitivity analyses to gain insight into model behaviour. By comparing different global sensitivity analysis (GSA) rankings and local sensitivity coefficients, we can identify key regulators of the metabolic pathway. Comparing the variance-based GSA rankings with the local sensitivity analysis, some differences present themselves, which reveal the extent to which coincident perturbations will have synergistic effects on system behaviour. Likewise, differences between first-order and total-effect sensitivities for other parameters reveal subtleties of interdependence among coordinated perturbations. For example, the V_max_ of ICD_c_ ranks low in the local analysis, but has a higher ranking in the total-effect GSA. The high total-effect ranking implies that the influence of ICD_c_ activity level on ICD_c_ flux is sensitive to the values of other parameters. In the local domain defined by our nominal model parameterization, the flux is robust to changes in ICD_c_ activity, but that robustness can be lost when other features of the system are perturbed.

### Sensitivities of model outputs

#### Pyruvate cycling ratio sensitivity rankings

Analyzing the sensitivity rankings in Figure 4 reveals that the V_max_ of PDH is the most influential parameter in terms of pyruvate cycling. However, comparison between the first-order and total-effect rankings (panels A and B) reveals that the V_max_’s of CIC, DIC, and MDH_m_ are the most significant in terms of modulating the dependence of pyruvate cycling on other parameter values (i.e. their first-order and total-effects are different). These three enzymes are known to be important participants in pyruvate cycling pathways [42].

#### NADPH sensitivity rankings

Referring to Figure 5, the total effect and local sensitivity rankings (panels A and D) indicate that the NADPH consumption rate has the most influence over the steady state NADPH concentration; this is consistent with the model structure. Next, comparing the first-order and total-effect rankings (panels A and B) we can draw two conclusions. First, because they are ranked high by both measures, the V_max_’s of pyruvate transporter (PYC) and DIC have significant impact on the NADPH concentration, regardless of perturbation to the other parameters. Second, the rankings of the V_max_’s of PDH, PC, and ME_c_ differ between first-order and total-effect measures, meaning that these parameters are influential in modulating the effect of pathway perturbations on NADPH concentration.

#### Cytosolic and mitochondrial pyruvate sensitivity rankings

The sensitivity ranking for the steady state cytosolic pyruvate concentration are shown in the supplementary text. They reveal that the V_max_’s of the pyruvate transporter (PYC) and citrate synthase (CS) have the most significant impact on pyruvate concentration. Moreover, the agreement between first-order and total-effect rankings implies that PYC and CS have a dominant effect on the pyruvate concentration, regardless of perturbations to other parts of the pathway.

#### PC flux sensitivity rankings

The sensitivity rankings of steady state PC flux, shown in the supplementary text, mirror the results in Figure 4. We find that V_max_’s of CIC, DIC, and MDH_m_ are the most significant in terms of modulating the dependence of PC flux on other parameter values (i.e., their first-order and total-effects are considerably different). However, the V_max_ of PDH has significant impact on the PC flux, which is consistent with the fact that these two reactions form a branch point.

#### Cytosolic isocitrate dehydrogenase sensitivity rankings

Recall that ICD_c_ is the main producer of NADPH. Considering sensitivities of the steady-state ICD_c_ flux (shown in the supplementary text: results similar to Figure 5), we find that the V_max_’s of CIC, DIC, PC and ME_c_ have the most influence over parameter interactions: their first-order rankings are low, while their total effect rankings are high. These parameters were also found to have significant influence over parameter dependence for steady state pyruvate concentration.

#### Effect of perturbation in PC V_max_ on the TCA and pyruvate cycling fluxes

As Figure 6 shows, perturbations in PC activity have their strongest effect on lactate dehydrogenase flux, which corroborates the finding that lactate concentration increases significantly in the case of PC knock-down [8]. Considering the response in the flux through ME_c_, PYC, DIC, and ICD_c_ to perturbations in PC activity, we see that the first-order and total-effect rankings are consistently high, indicating that the effect of PC perturbations on these fluxes is robust to changes in the values in other model parameters. These effects are compensatory: the local sensitivity coefficients are all negative (Figure 6D), indicating that an increased flux through PC is balanced by decreased flux through these enzymes. This corroborates the findings that perturbations to PC activity do not impact the pyruvate cycling ratio and that NADPH concentrations are not sensitive to perturbations in PC activity [8].

#### Model Predictions

To identify target points in the model that have a strong effect in modulating pyruvate cycling ratio and NADPH concentration, we compared sensitivity rankings of the parameters across all the sensitivity methods on the pyruvate cycling ratio and NADPH concentration. The sensitivity ranking shows that the V_max_’s of the pyruvate transporter (PYC) and citrate synthase (CS) have the most significant impact on both the pyruvate cycling ratio and the NADPH concentration across all the sensitivity measures. However, when we look at the rankings of the two different measures provided by the variance-based method: the total-effect and first-order rankings, we see that the rankings of V_max_’s of PYC and CS are similar across the model outputs, indicating that these strong effects are insensitive to perturbations in other parameters. In contrast, the V_max_’s of transport enzymes citrate isocitrate carrier (CIC) and dicarboxylate carrier (DIC), which also have significant impacts on NADPH concentration and pyruvate cycling ratio, show significant differences between first-order and total-effect rankings, and thus exert influence over the dependence on other model parameters. Knock-down of CIC and DIC has been shown to inhibit glucose stimulated insulin secretion [18]. Next, we compared the sensitivity rankings of cytosolic and mitochondrial pyruvate across all the sensitivity methods. Here, we again observe that the parameters discussed above exert strong influence on the pyruvate concentration. In addition, it has been shown that pyruvate entry into TCA cycle through PC flux is correlated with the GSIS [1]. Therefore, we compared sensitivities of the PC flux across all the sensitivity rankings to identify target candidates for perturbing the pyruvate cycling ratio. We observe that the PC flux is influenced significantly by the candidate targets CIC and DIC in both local and global sensitivity ranking.

Taking these results together, the model predicts that combined perturbations (e.g., of both PYC and CIC, or PYC and DIC) can have the double-effect of a significant impact on both NADPH concentration and pyruvate cycling ratio. These combined perturbations show the greatest promise as drug targets for modulation of the pyruvate cycling ratio and NADPH:NADP^+^ ratio and hence insulin secretion.

We simulated the model to predict the response to knock-down experiments. The model predicts that the knock-down of PYC alone has a significant impact on the pyruvate cycling ratio, but a lesser impact on NADPH concentration: when PYC activity is reduced by 10%, we see a 13% decrease in the pyruvate cycling ratio and a 5% decrease in NADPH concentration. In contrast, a 10% knock-down of DIC leads to a 12% drop in NADPH concentration and a .08% decrease in pyruvate cycling ratio. However, a combined knock-down of PYC and DIC by 10% each led to significant impact on both on the pyruvate cycling ratio and NADPH concentration: the pyruvate cycling ratio dropped by 30% and the NADPH concentration was reduced by 48%. This proposed combination knock-down could be addressed experimentally to provide further validation of the model and insight into GSIS.

This model is part of continuing effort to corroborate predictive models of ◻-cell metabolism with available data. This data has been collected from a range of cell types and using a range of assays; the resulting variability means that the goal of producing a fully predictive model of ◻-cell metabolism is a significant challenge. In our case, the model is unable to capture the effect on lactate concentration of ICD_c_ and PC knock-downs, and quantitative agreement with some pyruvate cycling data could not be attained [8]. These limitations suggest refinements of the model that could be made as more data becomes available.

The training and test studies described above were carried out on rat insulinoma-derived cell lines (INS-1). These cells exhibit a broad range of GSIS sensitivity and are thus useful for investigating the amplifying pathway response. More recent studies conducted on human islets have shown similar glucose-stimulated insulin release patterns [21], [43]. In the current study, we address the pyruvate cycling pathways using the INS-1 cell line data, because the data available from the human islet is insufficient for model training. The key pyruvate cycling enzymes ME_c_ and ICD_c_ exhibit the same high activity in both rat INS-1 cells and human islets. Furthermore, as in the rat cell line ME_m_ is abundant in human islet cells, underscoring the similarities of the important pyruvate cycling pathways. However, in comparison with the INS-1 cell line, human islets show reduced pyruvate carboxylase activity [43]. Previously, it has been reported that PC suppression might activate compensatory responses in β-cells either through increasing flux of downstream pyruvate cycling fluxes such as ICD_c_ or through a separate pathway involving acetyl-CoA [43]–[45]; these effects are not explicitly included in our model. As a further extension, the model could be expanded to describe the role of α-ketogulutarate in insulin regulation [46], [47].

## Acknowledgments

This work was supported by a Discovery Grant from the Canadian Natural Science and Engineering Research Council (NSERC).

## Abbreviations

GLC: glucose
F6P: fructose-6-phosphate
FBP: fructose-1,6-bisphosphate
GAP: glyceraldehyde 3-phosphate
DPG: 1,3-bisphospho-D-glycerate
PEP: phosphenol pyruvate
GT: glucose transporter
GK: glucokinase
PFK: 6-phosphofructokinase
FBA: fructose-bisphosphate aldolase
GAPD: glyceraldehyde 3-phosphate dehydrogenase
PGP: bisphosphoglyceratephosphatase
PK: pyruvate kinase
LDH: lactate dehydrogenase
ACO_m_: aconitase (mitochondrial)
ACO_c_: aconitase (cytosolic)
CIC: citrate carrier
DIC: dicarboxyrate carrier
CL_c_: citrate lyase (cytosolic)
CS: citrate synthase
FM: fumarase
IDH_m_: isocitrate dehydrogenase (mitochondrial)
IDH_c_: isocitrate dehydrogenase (NADP^+^) (cytosolic)
MDH_m_: malate dehydrogenase (mitochondrial)
MDH_c_: malate dehydrogenase (cytosolic)
OGC: oxoglutarate carrier
PC: pyruvate carboxylase
PDC: pyruvate dehydrogenase complex
PYC: pyruvate carrier
AKD: α-ketoglutarate dehydrogenase
SCS: succinyl-CoA synthetase
SDH: succinate dehydrogenase
ME_m_: malic enzyme (mitochondrial)
ME_c_: malic enzyme (cytosolic)

## Supplementary Text

### 1 Kinetic Mechanism Details

#### 1.1 Abbreviations

Subscripts c, m, and s denote cytoplasmic, mitochondrial, and transport enzymes or metabolites respectively. Species without subscripts are not separated into compartments.

**Table 1:**
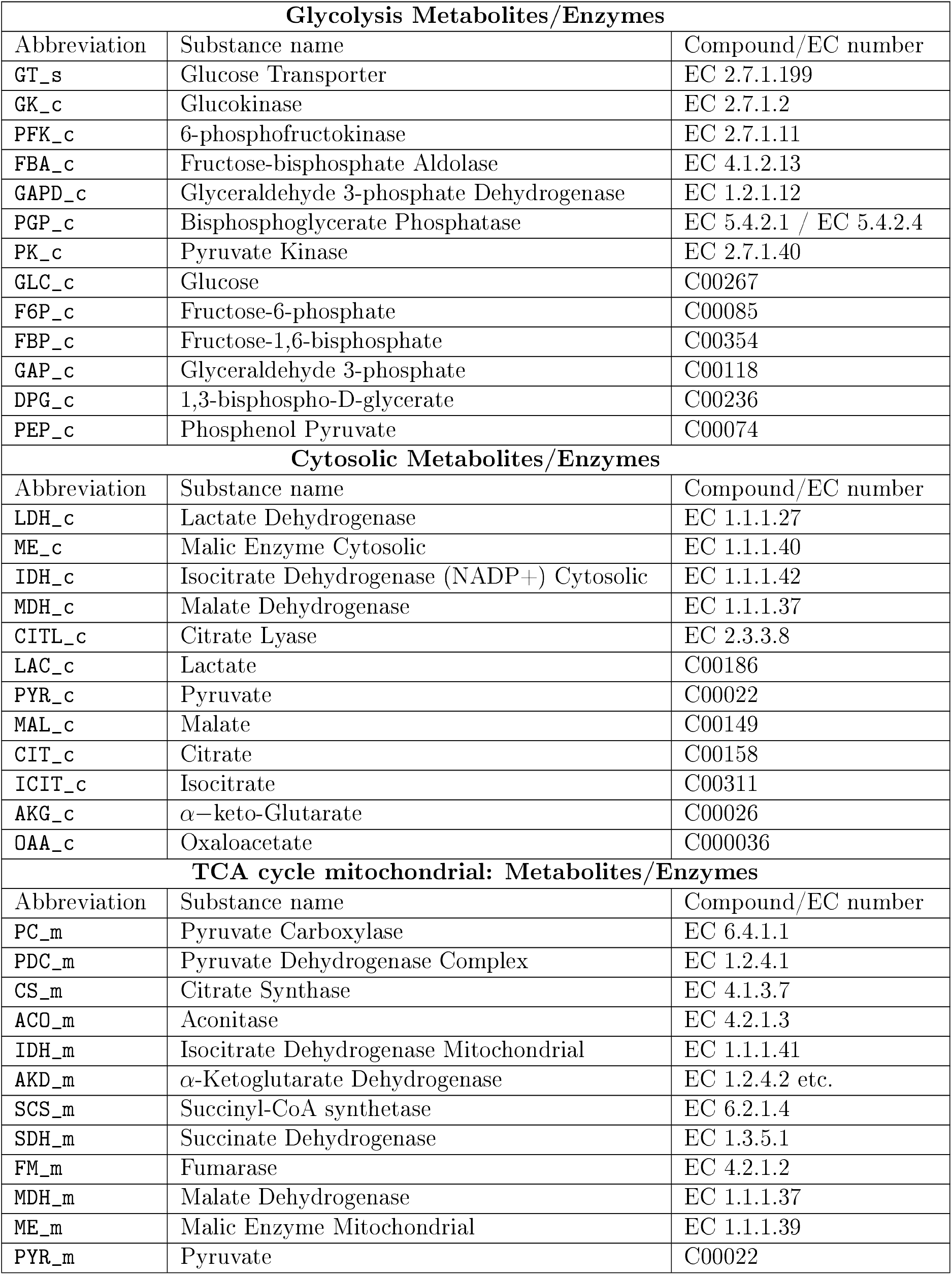
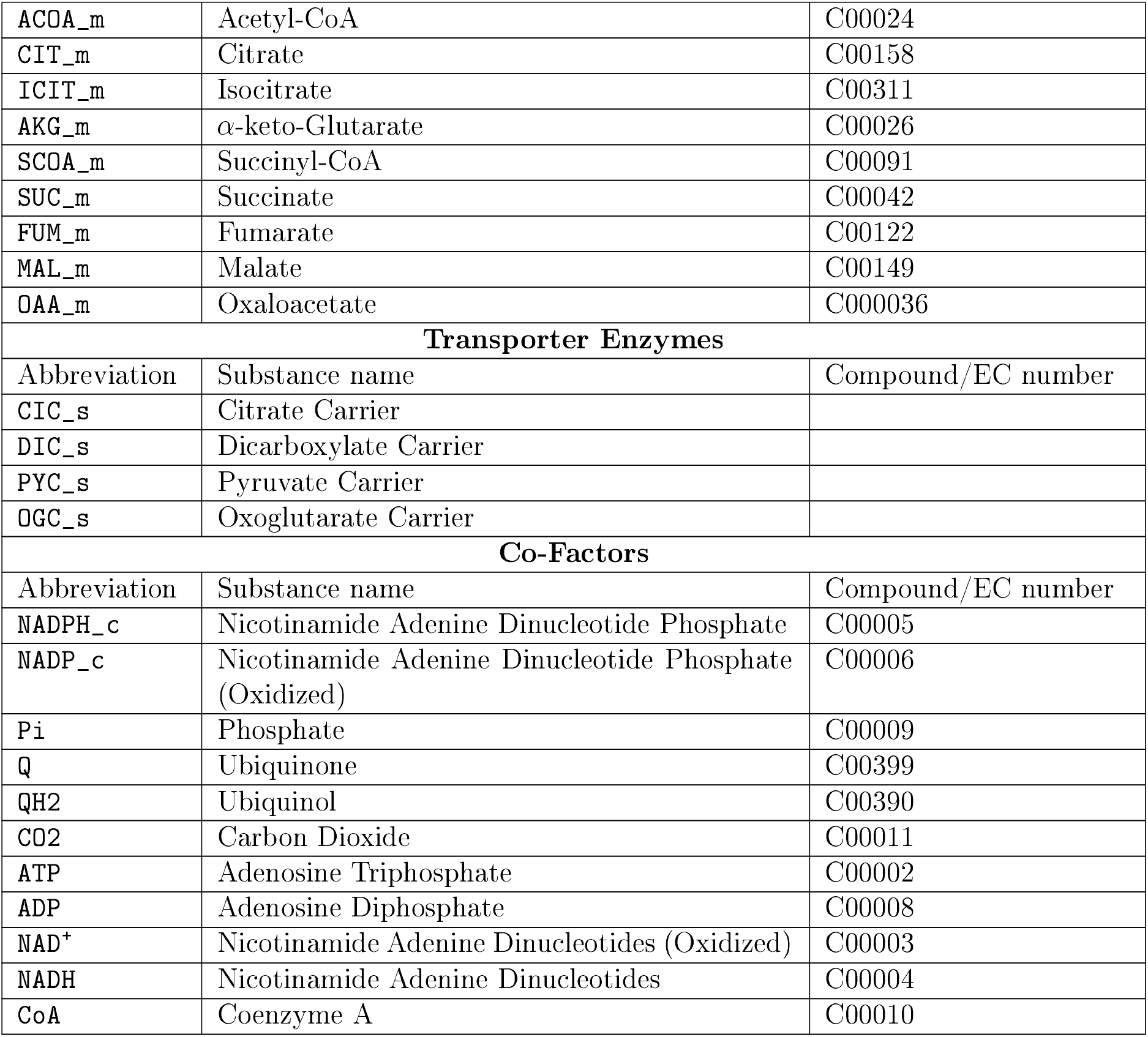
Abbreviations for Metabolites/Enzymes

| **Glycolysis Metabolites/Enzymes** |  |  |
| --- | --- | --- |
| Abbreviation | Substance name | Compound/EC number |
| GT_s | Glucose Transporter | EC 2.7.1.199 |
| GK_c | Glucokinase | EC 2.7.1.2 |
| PFK_c | 6-phosphofructokinase | EC 2.7.1.11 |
| FBA_c | Fructose-bisphosphate Aldolase | EC 4.1.2.13 |
| GAPD_c | Glyceraldehyde 3-phosphate Dehydrogenase | EC 1.2.1.12 |
| PGP_c | Bisphosphoglycerate Phosphatase | EC 5.4.2.1 / EC 5.4.2.4 |
| PK_c | Pyruvate Kinase | EC 2.7.1.40 |
| GLC_c | Glucose | C00267 |
| F6P_c | Fructose-6-phosphate | C00085 |
| FBP_c | Fructose-1,6-bisphosphate | C00354 |
| GAP_c | Glyceraldehyde 3-phosphate | C00118 |
| DPG_c | 1,3-bisphospho-D-glycerate | C00236 |
| PEP_c | Phosphenol Pyruvate | C00074 |
| **Cytosolic Metabolites/Enzymes** |  |  |
| Abbreviation | Substance name | Compound/EC number |
| LDH_c | Lactate Dehydrogenase | EC 1.1.1.27 |
| ME_c | Malic Enzyme Cytosolic | EC 1.1.1.40 |
| IDH_c | Isocitrate Dehydrogenase (NADP+) Cytosolic | EC 1.1.1.42 |
| MDH_c | Malate Dehydrogenase | EC 1.1.1.37 |
| CITL_c | Citrate Lyase | EC 2.3.3.8 |
| LAC_c | Lactate | C00186 |
| PYR_c | Pyruvate | C00022 |
| MAL_c | Malate | C00149 |
| CIT_c | Citrate | C00158 |
| ICIT_c | Isocitrate | C00311 |
| AKG_c | α-keto-Glutarate | C00026 |
| OAA_c | Oxaloacetate | C000036 |
| **TCA cycle mitochondrial: Metabolites/Enzymes** |  |  |
| Abbreviation | Substance name | Compound/EC number |
| PC_m | Pyruvate Carboxylase | EC 6.4.1.1 |
| PDC_m | Pyruvate Dehydrogenase Complex | EC 1.2.4.1 |
| CS_m | Citrate Synthase | EC 4.1.3.7 |
| ACO_m | Aconitase | EC 4.2.1.3 |
| IDH_m | Isocitrate Dehydrogenase Mitochondrial | EC 1.1.1.41 |
| AKD_m | α-Ketoglutarate Dehydrogenase | EC 1.2.4.2 etc. |
| SCS_m | Succinyl-CoA synthetase | EC 6.2.1.4 |
| SDH_m | Succinate Dehydrogenase | EC 1.3.5.1 |
| FM_m | Fumarase | EC 4.2.1.2 |
| MDH_m | Malate Dehydrogenase | EC 1.1.1.37 |
| ME_m | Malic Enzyme Mitochondrial | EC 1.1.1.39 |
| PYR_m | Pyruvate | C00022 |
| ACOA_m | Acetyl-CoA | C00024 |
| CIT_m | Citrate | C00158 |
| ICIT_m | Isocitrate | C00311 |
| AKG_m | α-keto-Glutarate | C00026 |
| SCOA_m | Succinyl-CoA | C00091 |
| SUC_m | Succinate | C00042 |
| FUM_m | Fumarate | C00122 |
| MAL_m | Malate | C00149 |
| OAA_m | Oxaloacetate | C000036 |
| **Transporter Enzymes** |  |  |
| Abbreviation | Substance name | Compound/EC number |
| CIC_s | Citrate Carrier |  |
| DIC_s | Dicarboxylate Carrier |  |
| PYC_s | Pyruvate Carrier |  |
| OGC_s | Oxoglutarate Carrier |  |
| **Co-Factors** |  |  |
| Abbreviation | Substance name | Compound/EC number |
| NADPH_c | Nicotinamide Adenine Dinucleotide Phosphate | C00005 |
| NADP_c | Nicotinamide Adenine Dinucleotide Phosphate (Oxidized) | C00006 |
| Pi | Phosphate | C00009 |
| Q | Ubiquinone | C00399 |
| QH2 | Ubiquinol | C00390 |
| CO2 | Carbon Dioxide | C00011 |
| ATP | Adenosine Triphosphate | C00002 |
| ADP | Adenosine Diphosphate | C00008 |
| NAD<sup>+</sup> | Nicotinamide Adenine Dinucleotides (Oxidized) | C00003 |
| NADH | Nicotinamide Adenine Dinucleotides | C00004 |
| CoA | Coenzyme A | C00010 |

#### 1.2 Model Reactions

**Table 2:**
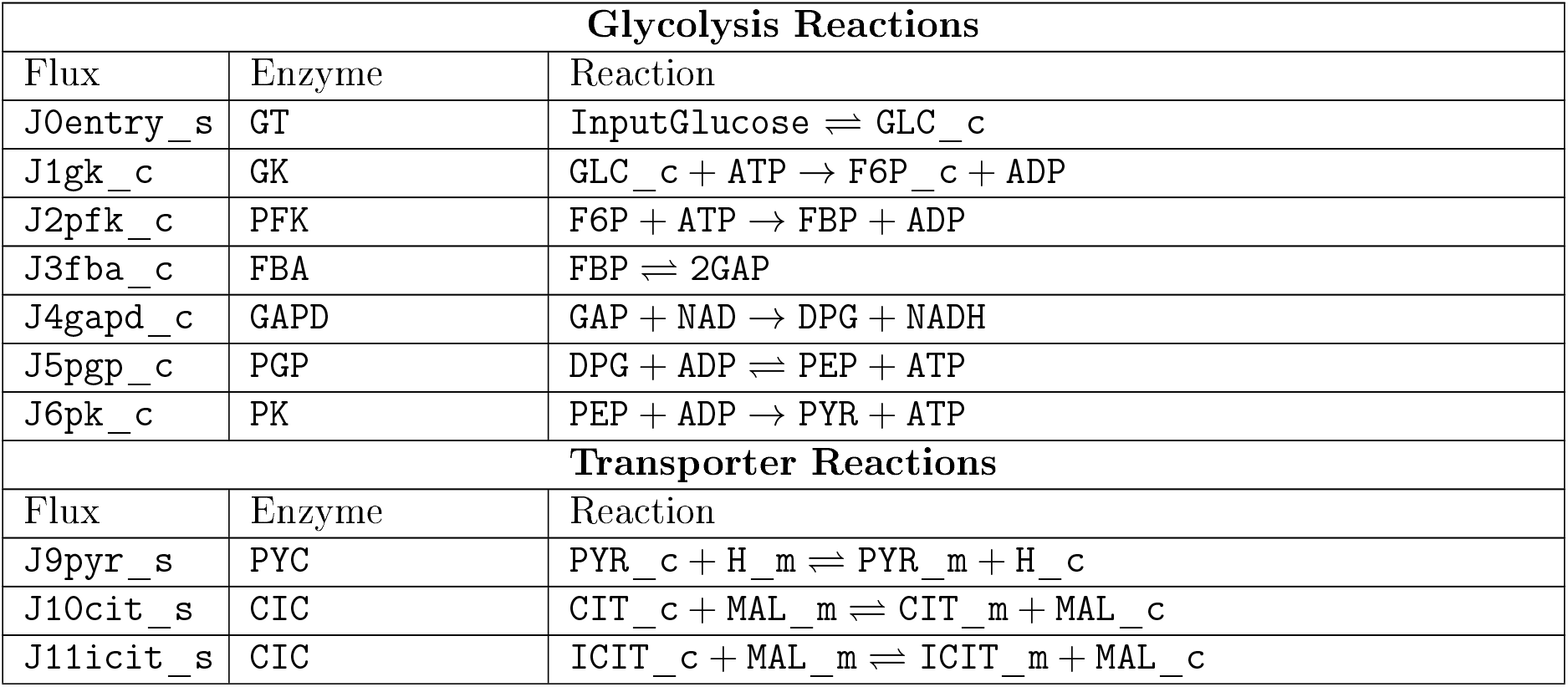
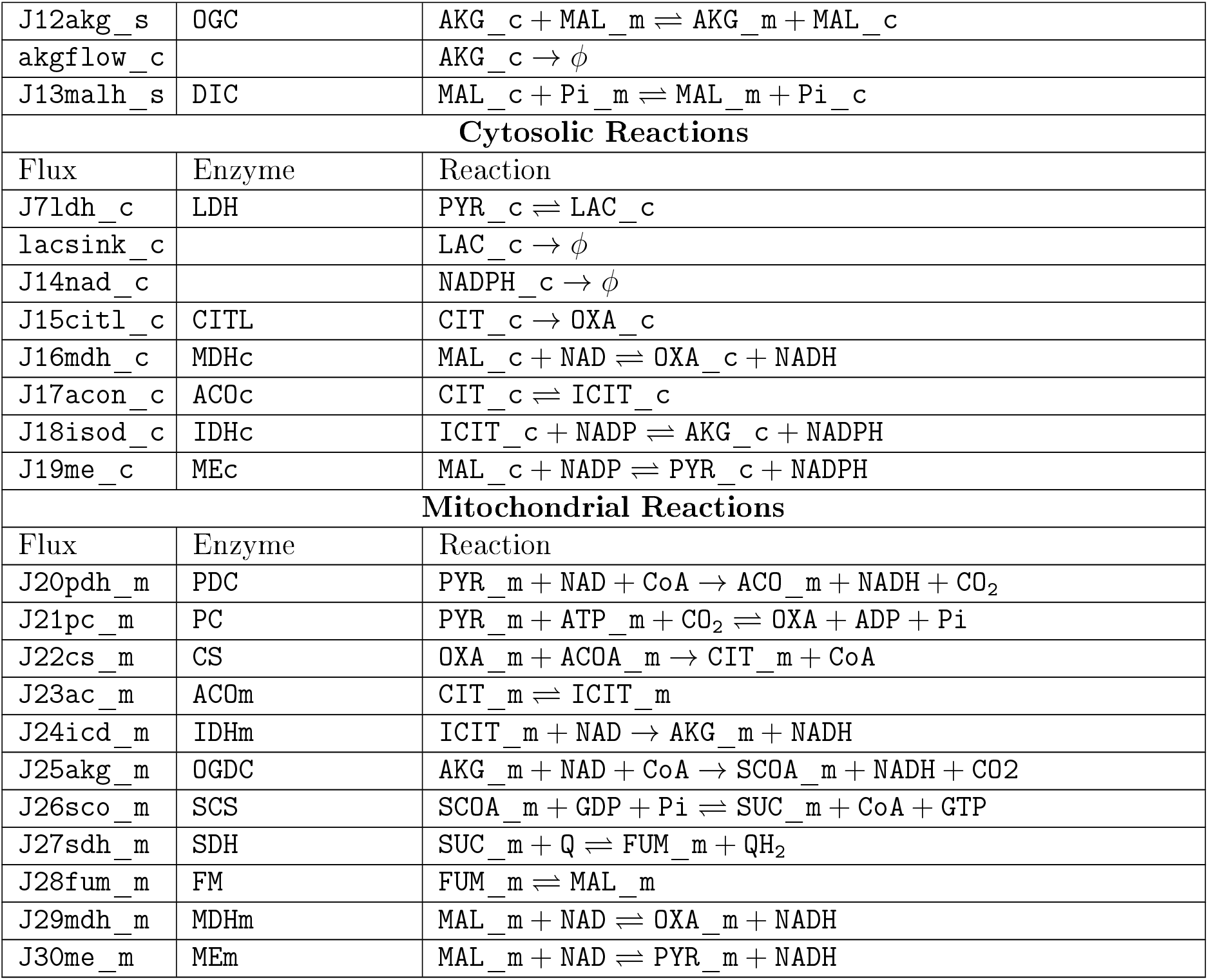
Model Reactions

| **Glycolysis Reactions** |  |  |
| --- | --- | --- |
| Flux | Enzyme | Reaction |
| J0entry_s | GT | $\text{InputGlucose} \rightleftharpoons \text{GLC\_c}$ |
| J1gk_c | GK | $\text{GLC\_c} + \text{ATP} \rightarrow \text{F6P\_c} + \text{ADP}$ |
| J2pfk_c | PFK | $\text{F6P} + \text{ATP} \rightarrow \text{FBP} + \text{ADP}$ |
| J3fba_c | FBA | $\text{FBP} \rightleftharpoons 2\text{GAP}$ |
| J4gapd_c | GAPD | $\text{GAP} + \text{NAD} \rightarrow \text{DPG} + \text{NADH}$ |
| J5pgp_c | PGP | $\text{DPG} + \text{ADP} \rightleftharpoons \text{PEP} + \text{ATP}$ |
| J6pk_c | PK | $\text{PEP} + \text{ADP} \rightarrow \text{PYR} + \text{ATP}$ |
| **Transporter Reactions** |  |  |
| Flux | Enzyme | Reaction |
| J9pyr_s | PYC | $\text{PYR\_c} + \text{H\_m} \rightleftharpoons \text{PYR\_m} + \text{H\_c}$ |
| J10cit_s | CIC | $\text{CIT\_c} + \text{MAL\_m} \rightleftharpoons \text{CIT\_m} + \text{MAL\_c}$ |
| J11icit_s | CIC | $\text{ICIT\_c} + \text{MAL\_m} \rightleftharpoons \text{ICIT\_m} + \text{MAL\_c}$ |
| J12akg_s | OGC | $AKG\_c + MAL\_m \rightleftharpoons AKG\_m + MAL\_c$ |
| akgflow_c | | $AKG\_c \rightarrow \phi$ |
| J13malh_s | DIC | $MAL\_c + Pi\_m \rightleftharpoons MAL\_m + Pi\_c$ |
| **Cytosolic Reactions** |  |  |
| Flux | Enzyme | Reaction |
| J7ldh_c | LDH | $PYR\_c \rightleftharpoons LAC\_c$ |
| lacsink_c | | $LAC\_c \rightarrow \phi$ |
| J14nad_c | | $NADPH\_c \rightarrow \phi$ |
| J15citl_c | CITL | $CIT\_c \rightarrow OXA\_c$ |
| J16mdh_c | MDHc | $MAL\_c + NAD \rightleftharpoons OXA\_c + NADH$ |
| J17acon_c | ACOc | $CIT\_c \rightleftharpoons ICIT\_c$ |
| J18isod_c | IDHc | $ICIT\_c + NADP \rightleftharpoons AKG\_c + NADPH$ |
| J19me_c | MEc | $MAL\_c + NADP \rightleftharpoons PYR\_c + NADPH$ |
| **Mitochondrial Reactions** |  |  |
| Flux | Enzyme | Reaction |
| J20pdh_m | PDC | $PYR\_m + NAD + CoA \rightarrow ACO\_m + NADH + CO_2$ |
| J21pc_m | PC | $PYR\_m + ATP\_m + CO_2 \rightleftharpoons OXA + ADP + Pi$ |
| J22cs_m | CS | $OXA\_m + ACOA\_m \rightarrow CIT\_m + CoA$ |
| J23ac_m | ACOm | $CIT\_m \rightleftharpoons ICIT\_m$ |
| J24icd_m | IDHm | $ICIT\_m + NAD \rightarrow AKG\_m + NADH$ |
| J25akg_m | OGDC | $AKG\_m + NAD + CoA \rightarrow SCoA\_m + NADH + CO_2$ |
| J26sco_m | SCS | $SCoA\_m + GDP + Pi \rightleftharpoons SUC\_m + CoA + GTP$ |
| J27sdh_m | SDH | $SUC\_m + Q \rightleftharpoons FUM\_m + QH_2$ |
| J28fum_m | FM | $FUM\_m \rightleftharpoons MAL\_m$ |
| J29mdh_m | MDHm | $MAL\_m + NAD \rightleftharpoons OXA\_m + NADH$ |
| J30me_m | MEm | $MAL\_m + NAD \rightleftharpoons PYR\_m + NADH$ |

#### 1.3 Fixed Species Concentrations and Biophysical Parameters

##### ATP and ADP

Cellular ATP and ADP concentrations are assumed to be directly dependent on the glucose concentration. This dependence is assumed to be piece-wise linear [1, 2, 3]. The ATP concentration is taken to increase linearly between 3mM and 7mM as glucose increases from 1mM to 10mM, with saturation at [ATP]= 7mM [2]. Similarly, the ADP concentration is taken to decrease linearly between 1.2mM and 0.6mM as glucose increases from 1mM to 10mM, with saturation at [ADP]= 0.6mM [2]. Finally, glucose serves as an input to the model and the input value is decided based on the simulation objective.

**Table 3:**
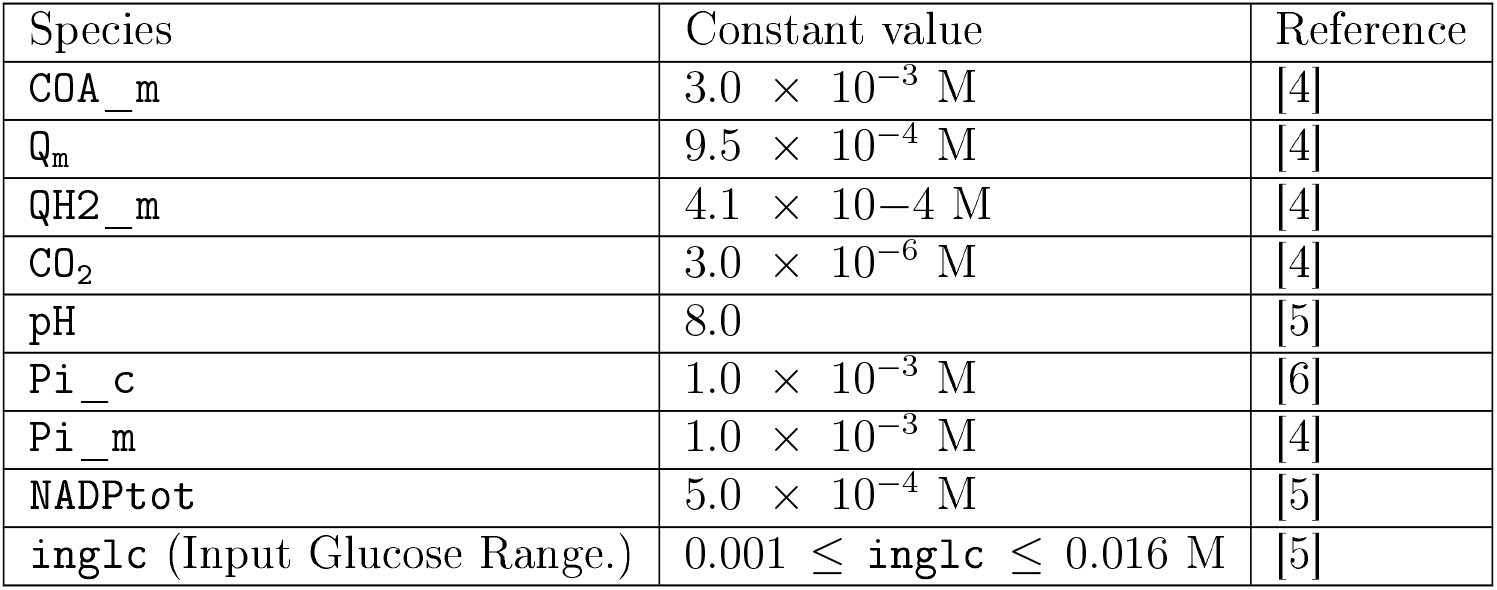
Fixed Values

#### 1.4 Rate Expressions

The model is divided in two parts: (i) the glycolysis model, and (ii) TCA cycle model, which includes the pyruvate cycling pathways. The glycolysis model is treated as an influx model for pyruvate cycling and was not included in the model analysis. The glycolysis model describes a six step pathway generating pyruvate, based on the work of Jiang et.al. [7]. The reaction kinetics and parameters are taken from SABIO-RK [8] Details are provided in Table 4.

Reaction kinetics for the enzymes in the TCA cycle model and the pyruvate cycling enzymes are reported in Table 4. These are adapted from Yugi and Tomita [9] and Westermark et.al. [10], with the exception of lactate dehydrogenase, which was adapted from Hoefnagel et.al. [11]. Furthermore, we modified the kinetics of cytosolic malic enzyme (from Westermark et.al. [10]) to incorporate NADPH as a dynamic variable; we used a rapid equilibrium random bi-bi mechanism expression to described the rate. Similarly, we modified the kinetics of mitochondrial malic enzyme (from Jiang et.al. [7]) to reversible Michaelis-Menten kinetics. The pyruvate dehydrogenase complex is subject to complex regulatory patterns involving Ca^2+^, NADH/NAD, Acetyl-coA, and phosphorylation-dephosphorylation [10]. It was not feasible to incorporate all these interactions. Instead, we simplified the kinetics from [10] to incorporate product inhibition by Acetyl-CoA. All the parameters for these modified kinetics are taken from the Brenda and SABIO-RK databases [8, 12]. Finally, for cytosolic isocitrate dehydrogenase we updated the parameters without modifying the kinetics from Brenda database [12]. We lumped NADPH consumption into a single reaction with rate described by irreversible Michaelis-Menten kinetics, with parameters estimated by fitting to the experimental data of Ronnebaum et. al. [5] as described in section 2.2.

To simplify the model description, we held the concentrations of ions and some cofactors listed in Table 1.3 as constant species. Rate expressions from the literature which involve these factors were simplified by defining effective rate constants that incorporate the fixed concentrations. Such simplifications were applied to the rate equations for the pyruvate transporter (`J9pyr_s`), the DIC carrier (`J13malh_s`), succinyl-coA synthetase (`J26sco_m`), and succinate dehydrogenase (`J27sdh_m`).

**Table 4:**
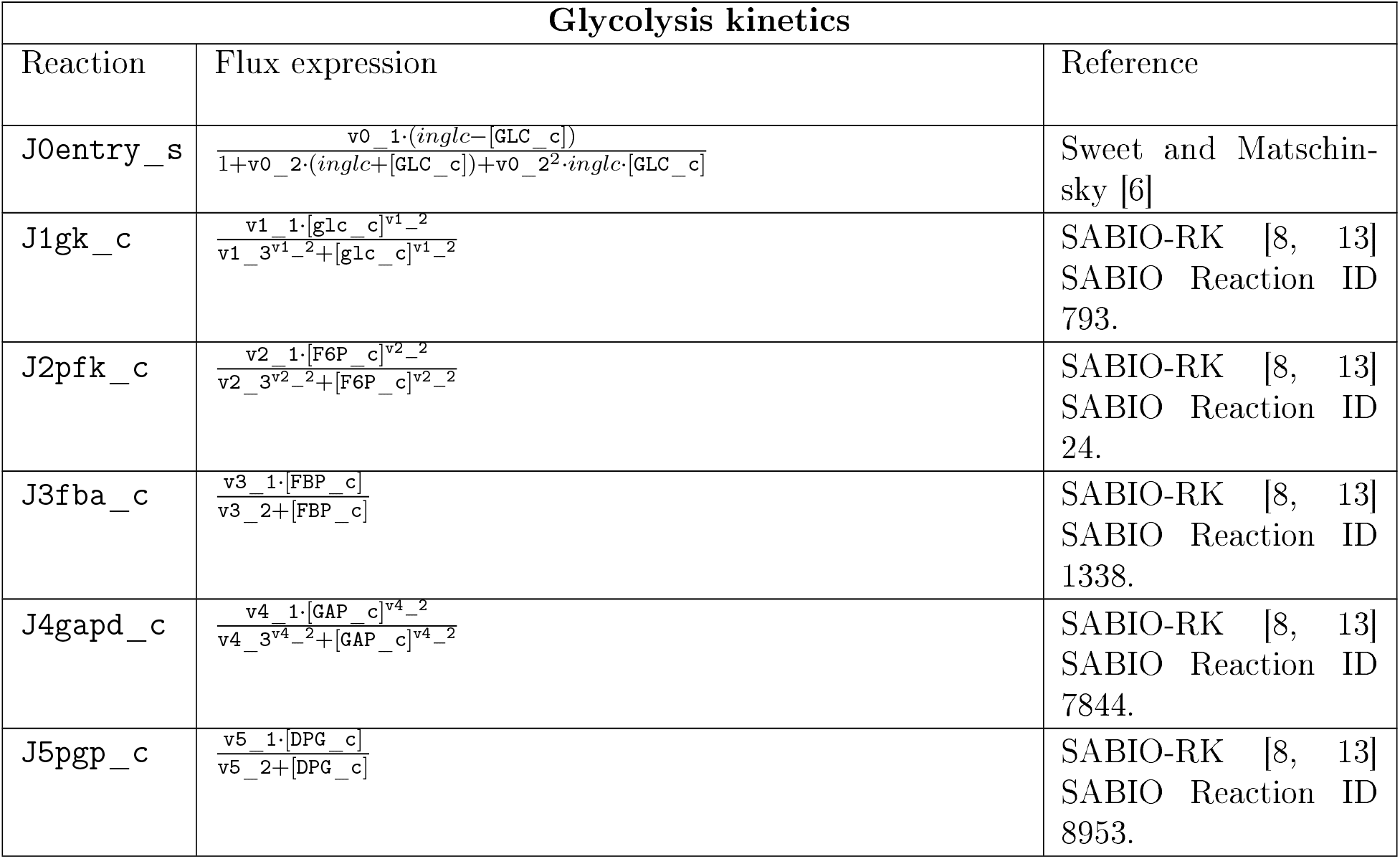
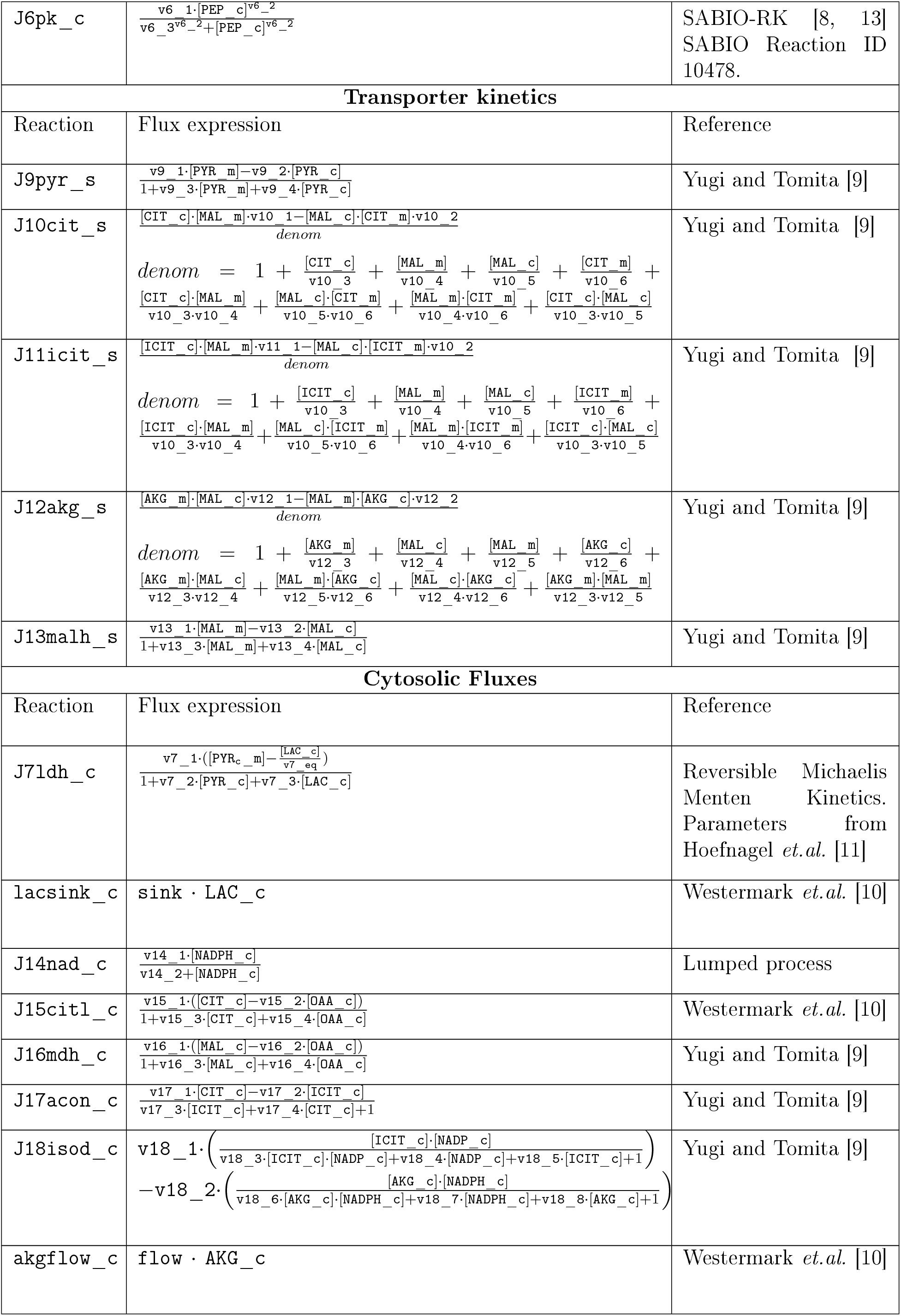
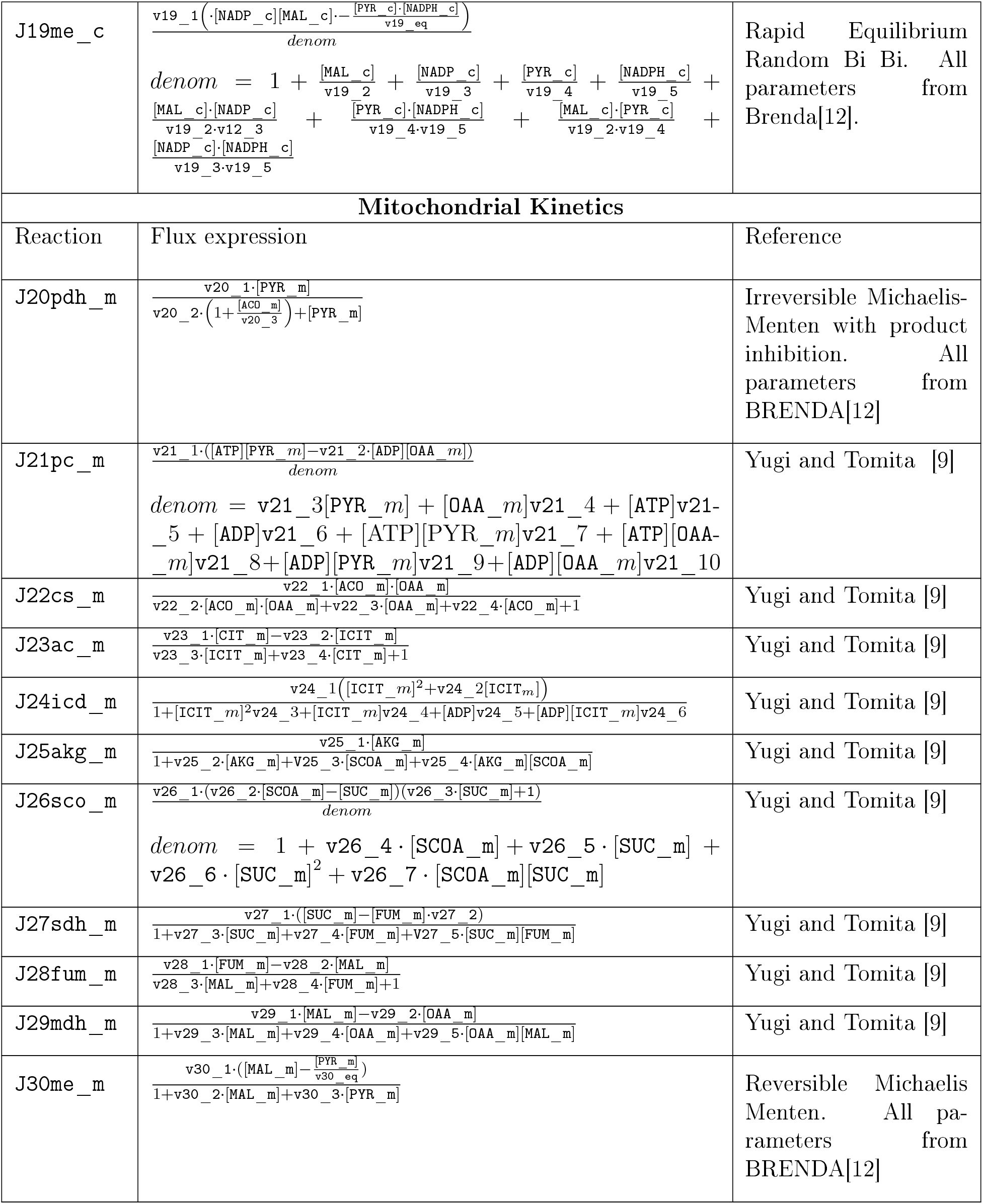
Kinetic Expressions

#### 1.5 Differential Equations

The model is described by the 24 differential equations. Next, NADP_c value is determined through conservation. Finally, ATP and ADP are modelled as a piecewise linear function.

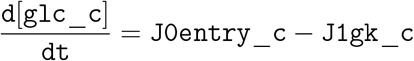

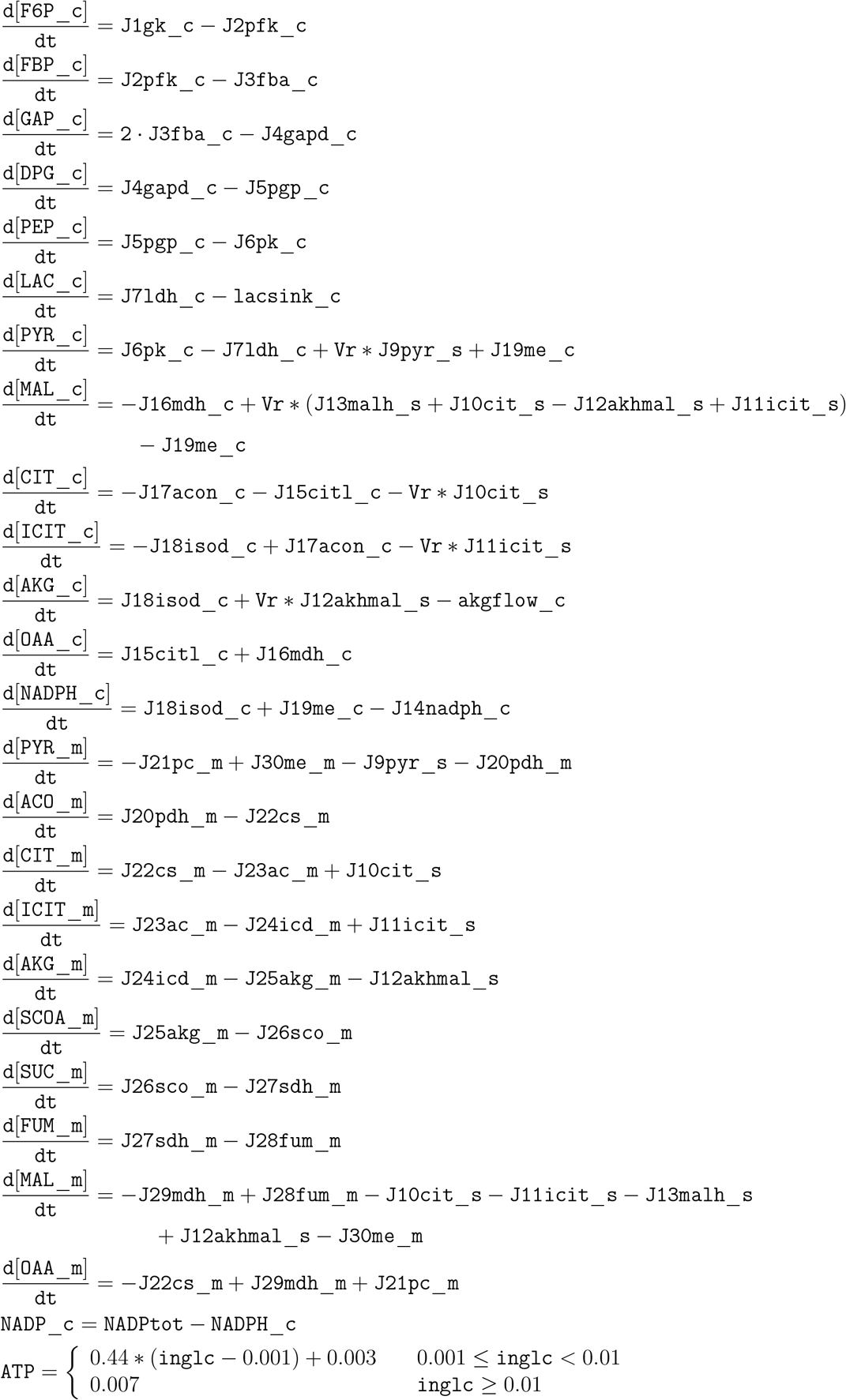

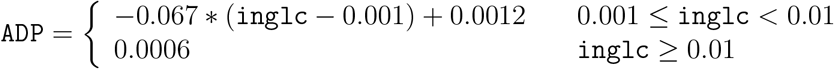

#### 1.6 Initial Conditions

The initial conditions for all simulations were obtained by integrating the system from the zero state for 4hrs (14400s) under appropriate glucose conditions.

### 2 Computational Settings for Solvers

#### 2.1 Steady State Calculation

To calculate the steady state we used the ode15s and fsolve functions of MATLAB^®^. The system of differential equations was first integrated up to 10^7^ seconds. The resulting state was then passed to fsolve to confirm the steady state conditions had been achieved. In order to increase stability and robustness of solvers, we generated a symbolic Jacobian using SBTOOLBOX2 [14, 15]. Since only 18% of the Jacobian coefficients were non-zero, we utilized the sparse storage mechanism as described in the ode15s and fsolve manual in order to increase the efficiency of solvers. We fixed the relative tolerance of ode15s at 10^−3^ and the absolute tolerance at 10^−6^, except for the states [F6Pj and [G6Pj for which we set the absolute tolerance at 10^−12^. For fsolve we used the default values of TolX and TolF (10^−6^).

#### 2.2 Parameter Optimization

The least-squares objective function for the parameter optimization is defined as.

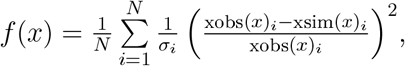

in which *N* is the number of experimental time-points, xobs(*x*)_*i*_ is the observed value of *i*^th^ state, xsim(*x*)_*i*_ is the corresponding *i*^th^ simulated state and *σ_i_* is the standard error of mean (SEM) of the *i*^th^ state. When the error was unknown then it was assumed to be 10%. All the state values are log scaled during optimization. The parameters were optimized in the range .01*p*_nom_ ≤ *p*_nom_ ≤ 100*p*_nom_, where *p*_nom_ is the nominal parameter value pulled from literature and databases. The optimization was carried out by a combination of the downhill simplex method in multiple dimensions and simulated annealing, implemented in the system biology toolbox [14].

#### 2.3 Global Sensitivity Analysis Settings

We used the variance-based Global Sensitivity Analysis (GSA, extended Fourier amplitude sensitivity test (eFAST) and the partial rank correlation method (PRCC) implemented in SBTOOLBOX2 [14, 15]. The analysis excluded the glycolysis influx mode, but treated all other model parameters. The relative parameter range variation was selected to be 100%. For the total sensitivity analysis, the objective function was defined as the sum of the squared errors between the observed and perturbed system output values:

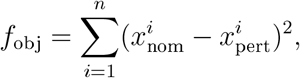

in which *x*_nom_ is the steady state nominal model output, *x*_pert_ is the steady state perturbed model output, and *n* is the number of state variables. In addition we calculated the sensitivities of individual state variables as a squared error:

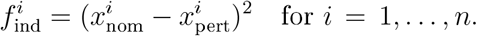

The pyruvate cycling rate is defined as the ratio of PC fiux and sum of the TCA cycle fiuxes [16]

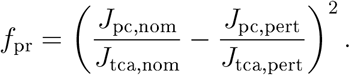

*f*_pr_ denotes the perturbation in the pyruvate cycling rate, *J*_pc_ is a fiux through the pyruvate carboxylase enzyme, and *J*_tca_ is a sum of all TCA cycle fluxes. Finally, nom and pert denote the nominal and perturbed values respectively.

The total number of model simulations was selected to be 10^5^, based on the suggestion of Saltelli [17] (2 × 512 × total number of parameters).

##### Integration Settings for Global Optimization

For simulating the model for global sensitivity analysis we used the SUNDIALS [18] package (MATLAB^®^ interface) in order to reduce the simulation time [14]. The final state was checked for steady-state as described previously using fsolve. The system was integrated using a relative tolerance of 10^−4^ and an absolute tolerance of 10^−14^ for all species concentrations. With these settings the output agreed with the MATLAB^®^ ode15s solver.

#### 2.4 Model Codes

The model is built in the MATLAB^®^ environment and the model codes are available at https://github.com/r2rahul/pyruvatecycling. The model is developed using SBToolbox2 [14, 15]. The code repository contains detail instructions on executing the model and reproducing the figures. Next, the model is available in the system biology markup language (SBML) format for the model interchange [19, 20, 21, 22, 23]

### 3 Parameters

The parameters were either pulled from the literature (Lit.) or estimated via the fitting exercise described above (Fit). For each parameter that was estimated, the reference from which the acceptable range was determined is listed. This model contains 129 global parameters. Units: Molarity (M) Seconds (s). Furthermore, Units are left blank for the dimensionless quantity.

**Table 5:** Model parameters.

| Parameter | Name (c:cytosol, m:mitochondria) | Value | Units | Fit or Lit. [References] |
| --- | --- | --- | --- | --- |
| v0_1 | Vmax (GLCc) | $2.7271 \cdot 10^{-6}$ | $s^{-1}$ | Fit [6] |
| v0_2 | Km (GLCc) | 101.0544 | $M^{-1}$ | Fit [6] |
| v1_1 | Vmax (GKc) | $1.3424 \cdot 10^{-4}$ | $M \cdot s^{-1}$ | Lit. [8, 13] |
| v1_2 | n (GKc) | $3.0118 \cdot 10^{-4}$ | | Lit. [8, 13] |
| v1_3 | Km (GKc) | 1.34 | M | Lit. [8, 13] |
| v2_1 | Vmax (PFKc) | $3.16667 \cdot 10^{-5}$ | $M \cdot s^{-1}$ | Lit. [8, 13] |
| v2_2 | n (PFKc) | 0.0089 |  | Lit. [8, 13] |
| v2_3 | Km (PFKc) | 0.9 | M | Lit. [8, 13] |
| v3_1 | Vmax (FBAc) | $7.97833 \cdot 10^{-5}$ | $M \cdot s^{-1}$ | Lit. [8, 13] |
| v3_2 | Km (FBAc) | $4 \cdot 10^{-6}$ | M | Lit. [8, 13] |
| v4_1 | Vmax (GAPc) | 0.001 | $M \cdot s^{-1}$ | Lit. [8, 13] |
| v4_2 | n (GAPc) | $3.2 \cdot 10^{-4}$ | | Lit. [8, 13] |
| v4_3 | Km (GAPc) | 1.5 | M | Lit. [8, 13] |
| v5_1 | Vmax (DPGc) | $3.33 \cdot 10^{-5}$ | $M \cdot s^{-1}$ | Lit. [8, 13] |
| v5_2 | Km (DPGc) | $8 \cdot 10^{-6}$ | M | Lit. [8, 13] |
| v6_1 | Vmax (PKc) | $5.33 \cdot 10^{-5}$ | $M \cdot s^{-1}$ | Lit. [8, 13] |
| v6_2 | n (PKc) | $1.5 \cdot 10^{-4}$ | | Lit. [8, 13] |
| v6_3 | Km (PKc) | 2.9 | M | Lit. [8, 13] |
| v7_1 | Vmax (LDHc) | 29.0969 | $s^{-1}$ | Fit [11] |
| v7_eq | Keq (LDHc) | 21.121 |  | Lit. [11] |
| v7_2 | Km1 (LDHc) | 448491.0 | $M^{-1}$ | Fit [11] |
| v7_3 | Km2(LDHc) | 449.0936 | $M^{-1}$ | Fit [11] |
| sink | Lactate Sink | $8.89 \cdot 10^{-8}$ | $M \cdot s^{-1}$ | Fit |
| v9_1 | Vmax(PYC, forward) | $3.7674 \cdot 10^{-8}$ | $s^{-1}$ | Lit. [9] |
| v9_2 | Vmax(PYC, reverse) | 0.004 | $s^{-1}$ | Fit [9] |
| v9_3 | Km1(PYC) | 49.2637 | $M^{-1}$ | Lit. [9] |
| v9_4 | Km2(PYC) | 187.3789 | $M^{-1}$ | Lit. [9] |
| v10_1 | Vmax(CIC, forward) | 32514.0 | $M^{-1} \cdot s^{-1}$ | Lit. [9] |
| v10_2 | Vmax(CIC, reverse) | 84267.0 | $M^{-1} \cdot s^{-1}$ | Lit. [9] |
| v10_3 | Km1(CIC) | $1.3 \cdot 10^{-4}$ | M | Lit. [9] |
| v10_4 | Km2(CIC) | $4.4 \cdot 10^{-4}$ | M | Lit. [9] |
| v10_5 | Km3(CIC) | $3.3 \cdot 10^{-4}$ | M | Lit. [9] |
| v10_6 | Km4(CIC) | $4.18 \cdot 10^{-5}$ | M | Lit. [9] |
| v12_1 | Vmax(OGC, forward) | 5811.9 | $M^{-1} \cdot s^{-1}$ | Lit. [9] |
| v12_2 | Vmax(OGC, reverse) | 6739.9 | $M^{-1} \cdot s^{-1}$ | Lit. [9] |
| v12_3 | Km1(OGC) | $3 \cdot 10^{-4}$ | M | Lit. [9] |
| v12_4 | Km2(OGC) | $7 \cdot 10^{-4}$ | M | Lit. [9] |
| v12_5 | Km3(OGC) | .0014 | M | Lit. [9] |
| v12_6 | Km4(OGC) | $1.7 \cdot 10^{-4}$ | M | Lit. [9] |
| v13_1 | Vmax(DIC, forward) | 0.8512 | $s^{-1}$ | Fit [9] |
| v13_2 | Vmax(DIC, reverse) | 0.2721 | $s^{-1}$ | Fit [9] |
| v13_3 | Km1(DIC) | 1388.9 | $M^{-1}$ | Lit. [9] |
| v13_4 | Vmax(DIC) | 1111.1 | $M^{-1}$ | Lit. [9] |
| v14_1 | Vmax(NADPH, sink) | 0.0025598 | $M \cdot s^{-1}$ | Fit |
| v14_2 | Km(NADPH, sink) | 0.98653 | M | Fit |
| v15_1 | Vmax(CITLc, forward) | $4.0008 \cdot 10^{-7}$ | $s^{-1}$ | Fit [10] |
| v15_2 | Vmax(CITLc, reverse) | $1.2 \cdot 10^{-4}$ | | Lit. [10] |
| v15_3 | Km1(CITLc) | $1.2 \cdot 10^{-4}$ | $M^{-1}$ | Lit. [10] |
| v15_4 | Km2(CITLc) | $1.2 \cdot 10^{-4}$ | $M^{-1}$ | Lit. [10] |
| v16_1 | Vmax(MDHc, forward) | 0.0036 | $s^{-1}$ | Lit. [9] |
| v16_2 | Vmax(MDHc, reverse) | 5000 | $s^{-1}$ | Lit. [9] |
| v16_3 | Km1(MDHc) | $8 \cdot 10^{-6}$ | $M^{-1}$ | Lit. [9] |
| v16_4 | Km2(MDHc) | 16667 | $M^{-1}$ | Lit. [9] |
| v17_1 | Vmax(ACONc, forward) | 0.0518 | $s^{-1}$ | Lit. [9] |
| v17_2 | Vmax(ACONc, forward) | 0.1104 | $s^{-1}$ | Fit [9] |
| v17_3 | Km2(ACONc) | 9090 | $M^{-1}$ | Lit. [9] |
| v17_4 | Km2(ACONc) | 2000 | $M^{-1}$ | Lit. [9] |
| v18_1 | Vmax(ISODc, forward) | 1152100 | $M^{-1} \cdot s^{-1}$ | Lit. [12] |
| v18_2 | Vmax(ISODc, reverse) | 5482500 | $M^{-1} \cdot s^{-1}$ | Lit. [12] |
| v18_3 | Km1(ISODc) | $2.3042 \cdot 10^{10}$ | $M^{-1}$ | Lit. [12] |
| v18_4 | Km2(ISODc) | 142857.1 | $M^{-1}$ | Fit [12] |
| v18_5 | Km3(ISODc) | 161290.3 | $M^{-1}$ | Lit. [12] |
| v18_6 | Km4(ISODc) | $1.0965 \cdot 10^{11}$ | $M^{-1}$ | Lit. [12] |
| v18_7 | Km5(ISODc) | 416666.7 | $M^{-1}$ | Lit. [12] |
| v18_8 | Km6(ISODc) | 263157.9 | $M^{-1}$ | Lit. [12] |
| flow | AKGc Sink | $1.46 \cdot 10^{-9}$ | $M \cdot s^{-1}$ | Fit |
| v19_1 | Vmax(MEc) | 913070 | $M^{-1} \cdot s^{-1}$ | Lit. [12] |
| v19_eq | Keq(MEc) | 1000 |  | Lit. [10] |
| v19_2 | Km1(MEc) | $1.2 \cdot 10^{-4}$ | M | Lit. [12] |
| v19_3 | Km2(MEc) | $1.39 \cdot 10^{-6}$ | M | Lit. [12] |
| v19_4 | Km3(MEc) | 0.0048 | M | Lit. [12] |
| v19_5 | Km4(MEc) | $5.3 \cdot 10^{-6}$ | M | Lit. [12] |
| v20_1 | Vmax(PDHm) | $1.417 \cdot 10^{-7}$ | $M \cdot s^{-1}$ | Fit [12] |
| v20_2 | Km(PDHm) | $3.5 \cdot 10^{-5}$ | M | Lit. [12] |
| v20_3 | Ki(PDHm) | $2 \cdot 10^{-5}$ | M | Lit. [12] |
| v21_1 | Vmax(PCm) | 0.1057 | $M^{-1} \cdot s^{-1}$ | Fit [9] |
| v21_2 | Keq(PCm) | 0.54 |  | Lit. [9] |
| v21_3 | Km1(PCm) | 4424100.0 | $M^{-1}$ | Lit. [9] |
| v21_4 | Km2(PCm) | $4.8528 \cdot 10^7$ | $M^{-1}$ | Lit. [9] |
| v21_5 | Km3(PCm) | 0.1057 | $M^{-1}$ | Fit [9] |
| v21_6 | Km4(PCm) | 0.54 | $M^{-1}$ | Lit. [9] |
| v21_7 | Km5(PCm) | 4424100.0 | $M^{-2}$ | Lit. [9] |
| v21_8 | Km6(PCm) | $4.8528 \cdot 10^7$ | $M^{-2}$ | Lit. [9] |
| v21_9 | Km7(PCm) | 0.1057 | $M^{-2}$ | Fit [9] |
| v21_10 | Km8(PCm) | 0.54 | $M^{-2}$ | Lit. [9] |
| v22_1 | Vmax(CSm) | 50.85 | $M^{-1} \cdot s^{-1}$ | Fit [9] |
| v22_2 | Km1(CSm) | $2.5 \cdot 10^{10}$ | $M^{-2}$ | Fit [9] |
| v22_3 | Km2(CSm) | 295000.0 | $M^{-1}$ | Lit. [9] |
| v22_4 | Km3(CSm) | 120000.0 | $M^{-1}$ | Fit [9] |
| v23_1 | Vmax(ACOm, forward) | 0.0518 | $s^{-1}$ | Lit. [9] |
| v23_2 | Vmax(ACOm, reverse) | 0.1104 | $s^{-1}$ | Lit. [9] |
| v23_3 | Km1(ACOm) | 9090.9 | $M^{-1}$ | Lit. [9] |
| v23_4 | Km2(ACOm) | 2000.0 | $M^{-1}$ | Lit. [9] |
| v24_1 | Vmax1(ICDm) | 0.1126 | $M^{-1} \cdot s^{-1}$ | Fit [9] |
| v24_2 | Vmax2(ICDm) | 0.0148 | M | Lit. [9] |
| v24_3 | Km1(ICDm) | 2777.5 | $M^{-2}$ | Lit. [9] |
| v24_4 | Km2(ICDm) | 0.63969 | $M^{-1}$ | Lit. [9] |
| v24_5 | Km3(ICDm) | 0.1126 | $M^{-2}$ | Fit [9] |
| v24_6 | Km4(ICDm) | 0.0148 | $M^{-2}$ | Lit. [9] |
| v25_1 | Vmax(AKGm) | 0.0311 | $s^{-1}$ | Fit [9] |
| v25_2 | Km1(AKGm) | $1.456 \cdot 10^9$ | $M^{-1}$ | Fit [9] |
| v25_3 | Km2(AKGm) | $1.4546 \cdot 10^9$ | $M^{-1}$ | Fit [9] |
| v25_4 | Km3(AKGm) | 24691.0 | $M^{-2}$ | Lit. [9] |
| v26_1 | Vmax(SCOm) | $3.32 \cdot 10^{-5}$ | $s^{-1}$ | Lit. [9] |
| v26_2 | Keq1(SCOm) | 6.4 |  | Lit. [9] |
| v26_3 | Ki(SCOm) | 4.1876 | $M^{-1}$ | Lit. [9] |
| v26_4 | Km1(SCOm) | 37.5348 | $M^{-1}$ | Lit. [9] |
| v26_5 | Km2(SCOm) | 1478.2 | $M^{-1}$ | Lit. [9] |
| v26_6 | Km3(SCOm) | 9509.6 | $M^{-2}$ | Lit. [9] |
| v26_7 | Km4(SCOm) | 236.7114 | $M^{-2}$ | Lit. [9] |
| v27_1 | Vmax(SDHm, forward) | 2.941 | $s^{-1}$ | Lit. [9] |
| v27_2 | Vmax(SDHm, reverse) | 11.5344 |  | Lit. [9] |
| v27_3 | Km1(SDHm) | 449.6613 | $M^{-1}$ | Lit. [9] |
| v27_4 | Km2(SDHm) | 199130.0 | $M^{-1}$ | Lit. [9] |
| v27_5 | Km3(SDHm) | $1.7921 \cdot 10^8$ | $M^{-2}$ | Lit. [9] |
| v28_1 | Vmax(FUMm, forward) | 13.9024 | $s^{-1}$ | Fit [9] |
| v28_2 | Vmax(FUMm, reverse) | 13.9024 | $s^{-1}$ | Lit. [9] |
| v28_3 | Km1(FUMm) | 4000000 | $M^{-1}$ | Lit. [9] |
| v28_4 | Km2(FUMm) | 880000 | $M^{-1}$ | Lit. [9] |
| v29_1 | Vmax(MDHm, forward) | .0016 | $s^{-1}$ | Lit. [9] |
| v29_2 | Vmax(MDHm, reverse) | $2.5563 \cdot 10^{-10}$ | $s^{-1}$ | Lit. [9] |
| v29_3 | Km1 (MDHm) | 33263.0 | $M^{-1}$ | Fit [9] |
| v29_4 | Km2 (MDHm) | 14.0754 | $M^{-1}$ | Lit. [9] |
| v29_5 | Km3 (MDHm) | 2742900 | $M^{-2}$ | Lit. [9] |
| v30_1 | Vmax (ME<sub>m</sub>) | 157.8125 | $s^{-1}$ | Fit [12] |
| v30_eq | Keq (ME<sub>m</sub>) | 1000 |  | Lit. [10] |
| v30_2 | Km1 (ME<sub>m</sub>) | 500 | $M^{-1}$ | Fit [12] |
| v30_3 | Km2 (ME<sub>m</sub>) | 227.2727 | $M^{-1}$ | Lit. [12] |
| Vr | Volume Ratio | 20 |  | Lit. [10] |

### 4 Supplementary Sensitivity Results

#### 4.1 Correlation Between States

To analyse the correlation between metabolite concentrations and the pyruvate cycling ratio, and between the metabolite concentrations and the NADPH level, we sampled the parameter space using latin hypercube sampling (LHS), recorded the output states and pyruvate cycling ratio, and calculated the correlations.

**Figure 1:**
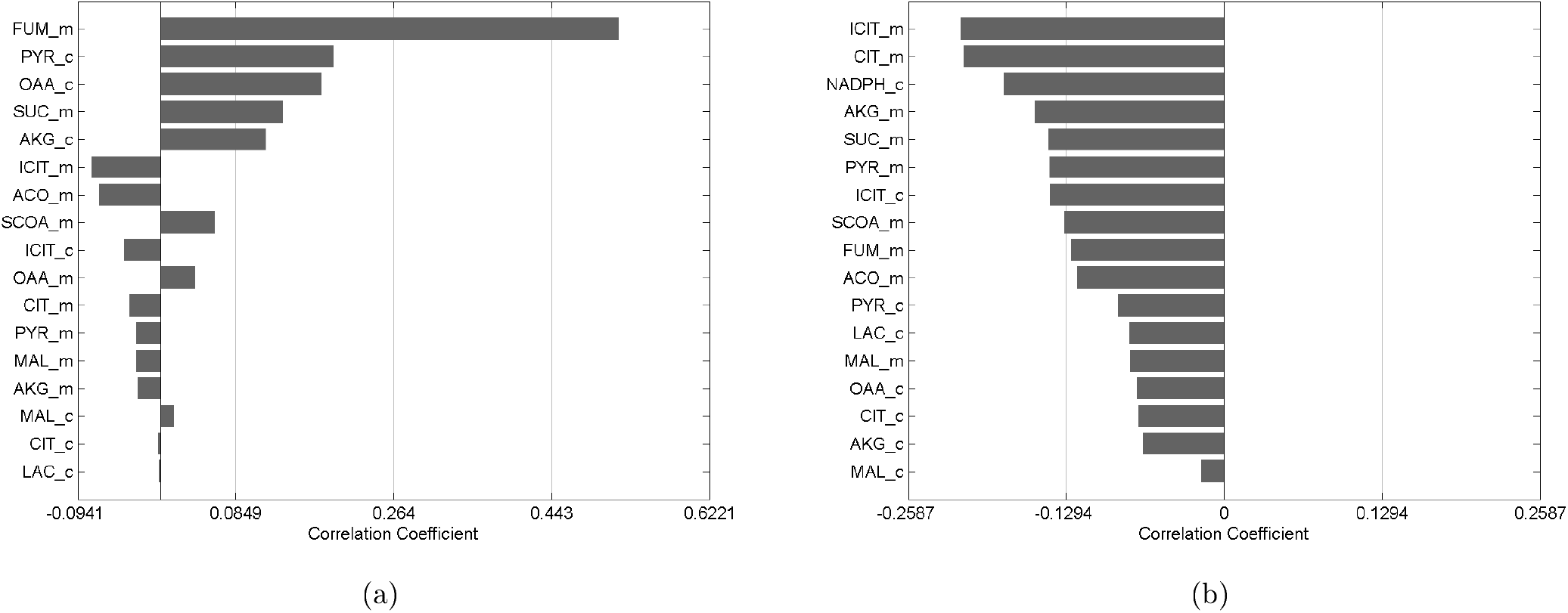
(a) Correlation between metabolite concentrations and the NADPH. (b) Correlation between metabolite concentrations and the pyruvate cycling ratio.

#### 4.2 Supplementary Sensitivity Figures

##### 4.2.1 Cytosolic Pyruvate Sensitivity Rankings

**Figure 2:**
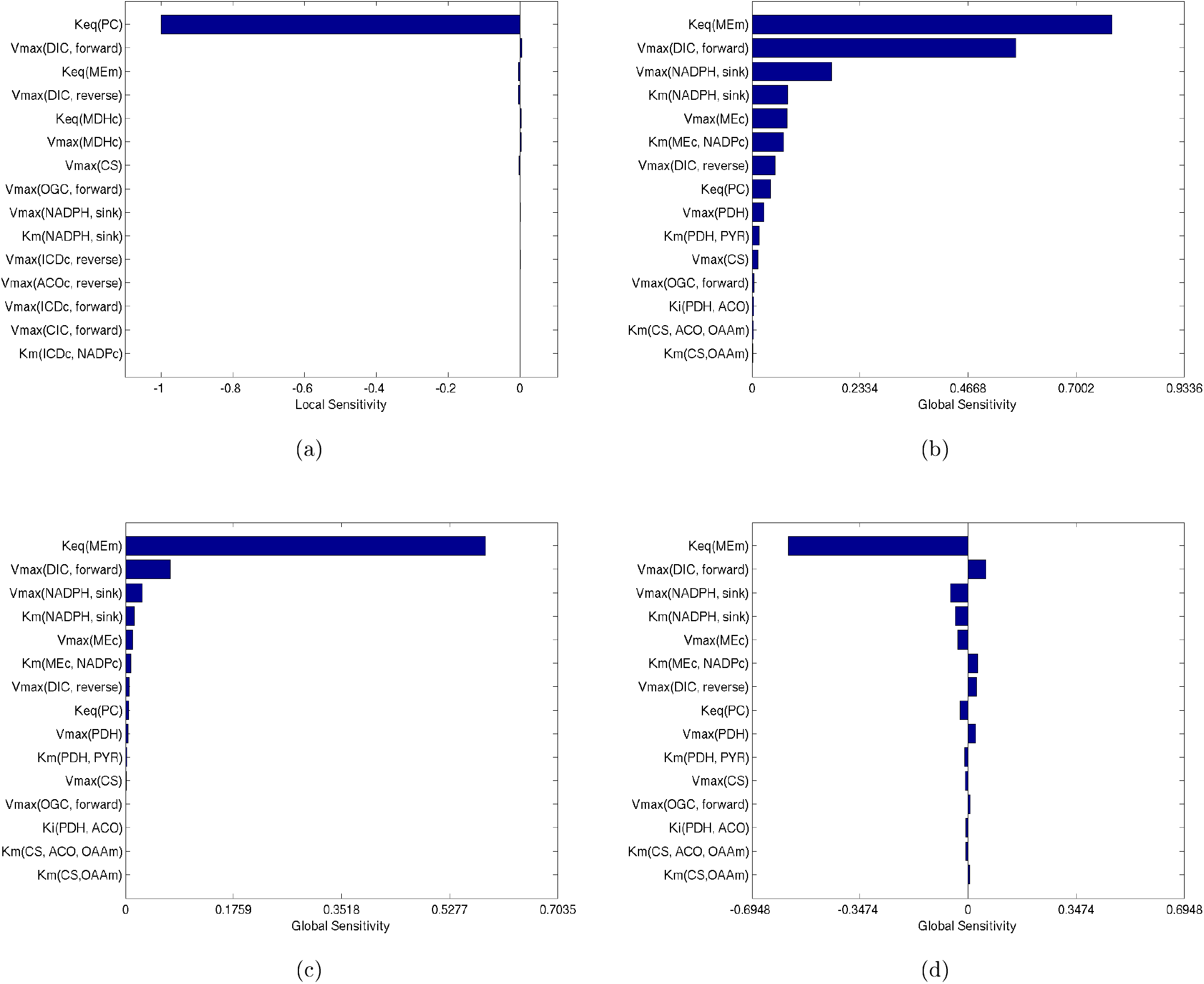
Global and local sensitivity rankings of cytosolic pyruvate concentration across different methods. (a) local sensitivity, (b) eFAST total effect, (c) eFAST first order, and (d) PRCC.

##### 4.2.2 Mitochondrial Pyruvate Sensitivity Rankings

**Figure 3:**
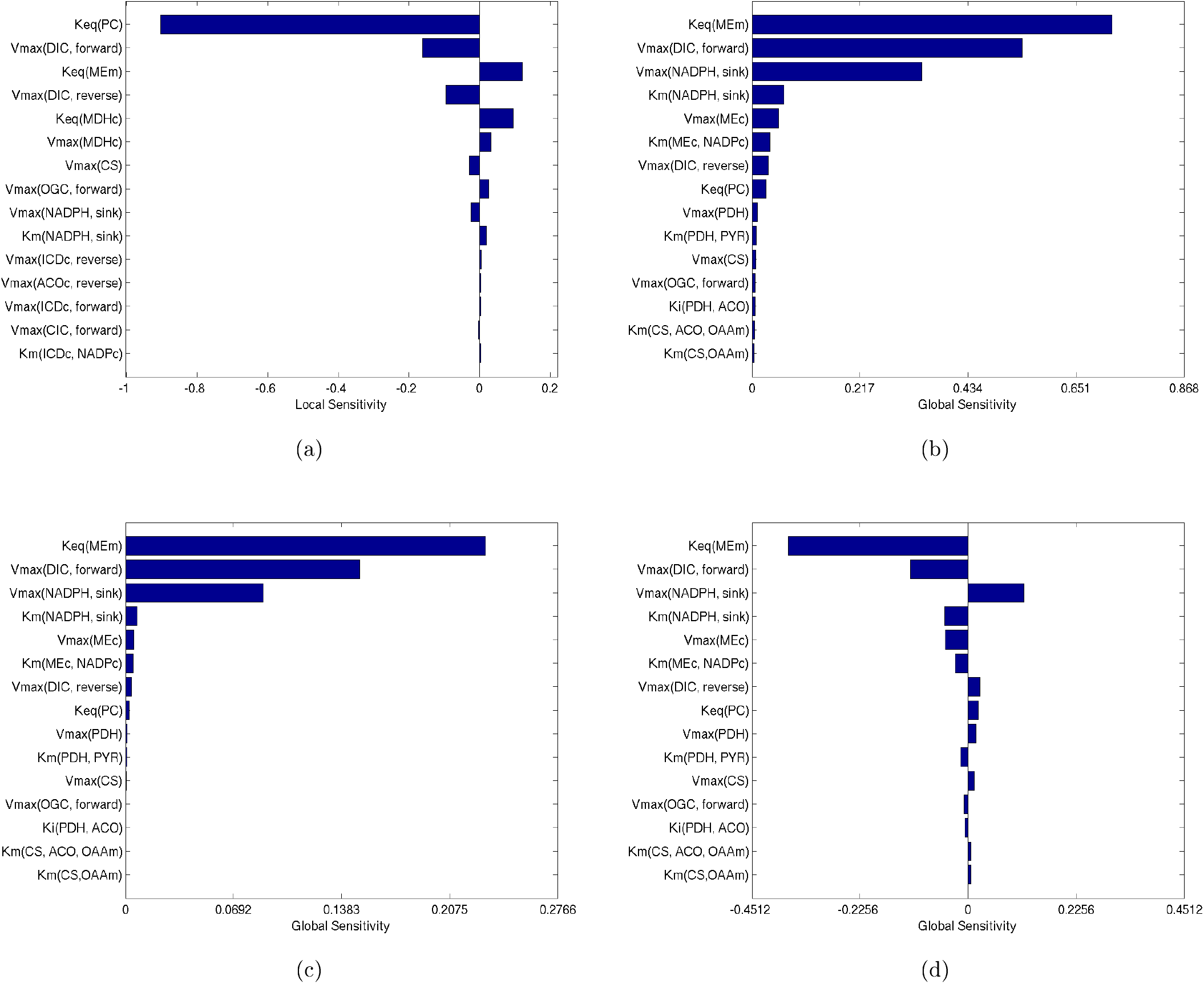
Global and local sensitivity rankings of mitochondrial pyruvate concentration across different methods. (a) local sensitivity, (b) eFAST total effect, (c) eFAST first order, and Panel (d) PRCC.

##### 4.2.3 Pyruvate Carboxylase Flux Sensitivity Rankings

**Figure 4:**
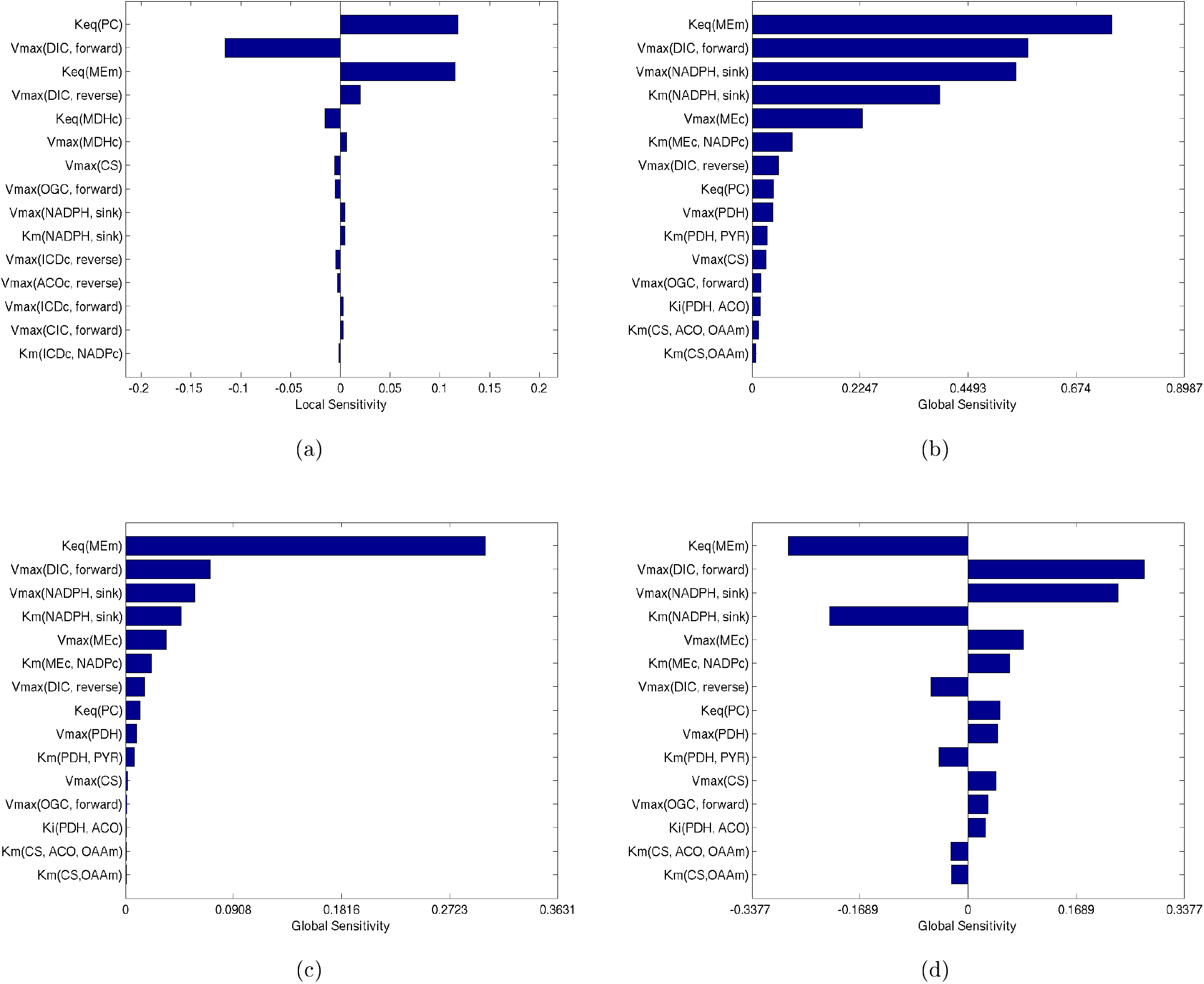
Global and local sensitivity rankings of pyruvate carboxylase flux across different methods. (a) local sensitivity, (b) eFAST total effect, (c) eFAST first order, and (d) PRCC.

##### 4.2.4 Cytosolic Isocitrate Dehydrogenase Flux Sensitivity Rankings

**Figure S5:**
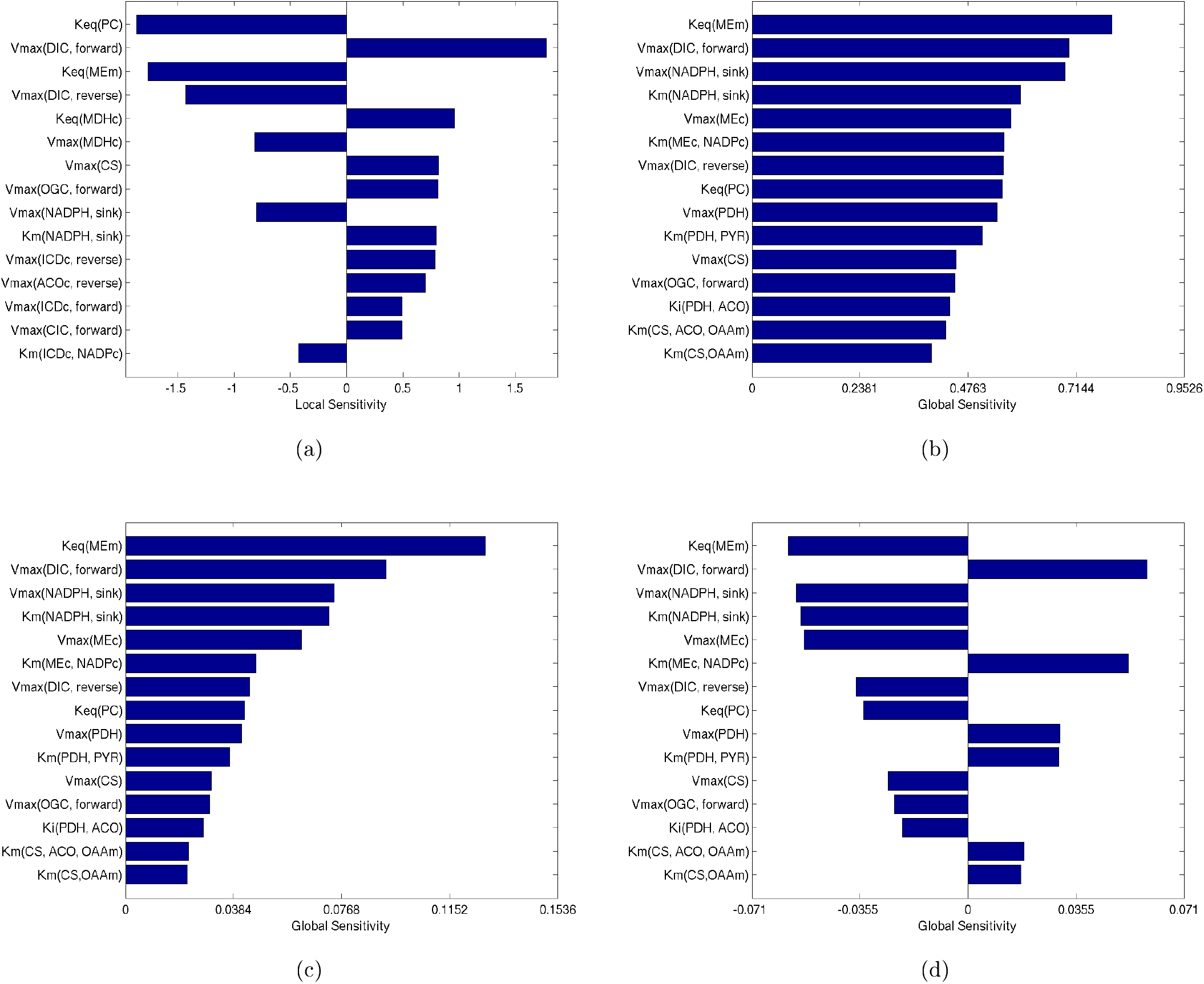
Global and local sensitivity rankings of cytosolic isocitrate dehydrogenase flux across different methods. (a) local sensitivity, (b) eFAST total effect, (c) eFAST first order, and (d) PRCC.

## References

[1] D. M. Muoio and C. B. Newgard, “Mechanisms of disease: molecular and metabolic mechanisms of insulin resistance and beta-cell failure in type 2 diabetes.,” Nat. Rev. Mol. Cell Biol., vol. 9, no. 3, pp. 193–205, Mar. 2008.

[2] C. B. Newgard and F. M. Matschinsky, “Handbook of Physiology.” Nature Publishing Group, 2001.

[3] J. C. Henquin, M. a Ravier, M. Nenquin, J. C. Jonas, and P. Gilon, “Hierarchy of the beta-cell signals controlling insulin secretion.,” Eur. J. Clin. Invest., vol. 33, no. 9, pp. 742–50, Sep. 2003.

[4] J. C. Henquin, “Triggering and amplifying pathways of regulation of insulin secretion by glucose.,” Diabetes, vol. 49, no. 11, pp. 1751–60, Nov. 2000.

[5] P. O. Westermark, J. H. Kotaleski, A. Björklund, V. Grill, and A. Lansner, “A mathematical model of the mitochondrial NADH shuttles and anaplerosis in the pancreatic beta-cell.,” Am. J. Physiol. Endocrinol. Metab., vol. 292, no. 2, pp. E373–93, Feb. 2007.

[6] K. Yugi and M. Tomita, “A general computational model of mitochondrial metabolism in a whole organelle scale.,” Bioinformatics, vol. 20, no. 11, pp. 1795–6, Jul. 2004.

[7] Z.-H. Yang, J.-G. Liu, C.-R. Yu, and J.-T. Han, “Quantifying the effect of investors’ attention on stock market,” PLoS One, vol. 12, no. 5, p. e0176836, May 2017.

[8] S. M. Ronnebaum et al., “A pyruvate cycling pathway involving cytosolic NADP-dependent isocitrate dehydrogenase regulates glucose-stimulated insulin secretion.,” J. Biol. Chem., vol. 281, no. 41, pp. 30593–602, Oct. 2006.

[9] N. Sekine et al., “Low Lactate Dehydrogenase and High Mitochondrial Glycerol Phosphate Dehydrogenase in Pancreatic P-Cells,” vol. 269, no. 7, pp. 4895–4902, 1994.

[10] C. J. Hedeskov, K. Capito, and P. Thams, “Cytosolic ratios of free [NADPH]/[NADP+] and [NADH]/[NAD+] in mouse pancreatic islets, and nutrient-induced insulin secretion.,” Biochem. J., vol. 241, no. 1, pp. 161–7, Jan. 1987.

[11] A Sener, F. Malaisse-Lagae, S. P. Dufrane, and W. J. Malaisse, “The coupling of metabolic to secretory events in pancreatic islets. The cytosolic redox state.,” Biochem. J., vol. 220, no. 2, pp. 433–40, Jun. 1984.

[12] J. W. Joseph et al., “Normal flux through ATP-citrate lyase or fatty acid synthase is not required for glucose-stimulated insulin secretion.,” J. Biol. Chem., vol. 282, no. 43, pp. 31592–600, Oct. 2007.

[13] M. V Jensen et al., “Compensatory responses to pyruvate carboxylase suppression in islet beta-cells. Preservation of glucose-stimulated insulin secretion.,” J. Biol. Chem., vol. 281, no. 31, pp. 22342–51, Aug. 2006.

[14] S. M. Ronnebaum et al., “Silencing of cytosolic or mitochondrial isoforms of malic enzyme has no effect on glucose-stimulated insulin secretion from rodent islets.,” J. Biol. Chem., vol. 283, no. 43, pp. 28909–17, Oct. 2008.

[15] M. J. MacDonald, “Feasibility of a mitochondrial pyruvate malate shuttle in pancreatic islets. Further implication of cytosolic NADPH in insulin secretion,” J. Biol. Chem., vol. 270, no. 34, pp. 20051–20058, Aug. 1995.

[16] D. Lu et al., “13C NMR isotopomer analysis reveals a connection between pyruvate cycling and glucose-stimulated insulin secretion (GSIS).,” Proc. Natl. Acad. Sci. U. S. A., vol. 99, no. 5, pp. 2708–13, Mar. 2002.

[17] J. Xu et al., “Malic enzyme is present in mouse islets and modulates insulin secretion.,” Diabetologia, vol. 51, no. 12, pp. 2281–9, Dec. 2008.

[18] J. W. Joseph et al., “The mitochondrial citrate/isocitrate carrier plays a regulatory role in glucose-stimulated insulin secretion.,” J. Biol. Chem., vol. 281, no. 47, pp. 35624–32, Nov. 2006.

[19] R. G. Kibbey, R. L. Pongratz, A. J. Romanelli, C. B. Wollheim, G. W. Cline, and G. I. Shulman, “Mitochondrial GTP regulates glucose-stimulated insulin secretion.,” Cell Metab., vol. 5, no. 4, pp. 253–64, Apr. 2007.

[20] M. E. Rabaglia et al., “alpha-Ketoisocaproate-induced hypersecretion of insulin by islets from diabetes-susceptible mice,” Lloydia (Cincinnati), vol. 53706, pp. 218–224, 2005.

[21] M. J. MacDonald, M. J. Longacre, and M. A. Kendrick, “Mitochondrial malic enzyme (ME2) in pancreatic islets of the human, rat and mouse and clonal insulinoma cells: Simple enzyme assay for mitochondrial malic enzyme 2,” Arch. Biochem. Biophys., vol. 488, no. 2, pp. 100–104, 2009.

[22] P. E. MacDonald, J. W. Joseph, and P. Rorsman, “Glucose-sensing mechanisms in pancreatic beta-cells.,” Philos. Trans. R. Soc. Lond. B. Biol. Sci., vol. 360, no. 1464, pp. 2211–25, Dec. 2005.

[23] K. Eto et al., “Role of NADH Shuttle System in Glucose-Induced Activation of Mitochondrial Metabolism and Insulin Secretion,” Science (80-.)., vol. 283, no. 5404, pp. 981–985, Feb. 1999.

[24] N. Jiang, R. D. Cox, and J. M. Hancock, “A kinetic core model of the glucose-stimulated insulin secretion network of pancreatic β cells,” Mamm. Genome, vol. 18, no. 6-7, pp. 508–520, Jul. 2007.

[25] U. Wittig, M. Rey, A. Weidemann, R. Kania, and W. M. Üller, “SABIO-RK: an updated resource for manually curated biochemical reaction kinetics,” Nucleic Acids Res., vol. 46, 2018.

[26] U. Wittig et al., “SABIO-RK database for biochemical reaction kinetics,” Nucleic Acids Res., vol. 40, no. D1, pp. D790–D796, 2012.

[27] F. M. Sweet, I. R. and Matschinsky, “Mathematical model of beta-cell glucose metabolism and insulin release. I. Glucokinase as glucosensor hypothesis,” Am. J. Physiol. - Endocrinol. Metab., vol. 268, no. 4, pp. E775–E788, 1995.

[28] A. Chang et al., “BRENDA, the ELIXIR core data resource in 2021: new developments and updates,” Nucleic Acids Res., vol. 49, 2021.

[29] M. Scheer et al., “BRENDA, the enzyme information system in 2011,” Nucleic Acids Res., vol. 39, no. Database issue, pp. D670–676, Jan. 2011.

[30] H. Schmidt and M. Jirstrand, “Systems Biology Toolbox for MATLAB: a computational platform for research in systems biology,” Bioinformatics, vol. 22, no. 4, pp. 514–515, Feb. 2006.

[31] K. Tiwari et al., “Reproducibility in systems biology modelling,” Mol. Syst. Biol., vol. 17, no. 2, p. e9982, 2021.

[32] R. S. Malik-Sheriff et al., “BioModels-15 years of sharing computational models in life science,” Nucleic Acids Res., vol. 48, pp. 407–415, 2019.

[33] S. M. Keating et al., “SBML Level 3: an extensible format for the exchange and reuse of biological models,” Mol. Syst. Biol., vol. 16, no. 8, p. e9110, Aug. 2020.

[34] L. Eldén, L. Wittmeyer-Koch, and H. B. Nielson, Introduction to Numerical Computation: Analysis and MATLAB Illustrations. Lund, Sweden: Studentlitteratur AB, 2004.

[35] Y. Zheng, A. Rundell, and Y. Zhang, “Comparative study of parameter sensitivity analyses of the TCR-activated Erk-MAPK signalling pathway.,” Syst. Biol. (Stevenage)., vol. 153, no. 4, pp. 201–11, Jul. 2006.

[36] N. R. Draper and H. Smith, Applied regression analysis. Wiley, 1998.

[37] A. Saltelli, S. Tarantola, and K. P.-S. Chan, “A Quantitative Model-Independent Method for Global Sensitivity Analysis of Model Output,” Technometrics, vol. 41, no. 1, pp. 39–56, Feb. 1999.

[38] A. Saltelli, Global Sensitivity Analysis: The Primer. John Wiley, 2008.

[39] F. Campolongo, S. Tarantola, and a. Saltelli, “Sensitivity Anaysis as an Ingredient of Modeling,” Stat. Sci., vol. 15, no. 4, pp. 377–395, Nov. 2000.

[40] H. Schmidt, “IQM Tools Repository,” Iqmtools.intiquan.com. 2021.

[41] A. Sener and W. J. Malaisse, “The coupling of metabolic to secretory events in pancreatic islets: comparison between insulin release and cytosolic redox state.,” Biochem. Int., vol. 14, no. 5, pp. 897–902, May 1987.

[42] M. J. MacDonald, L. a Fahien, L. J. Brown, N. M. Hasan, J. D. Buss, and M. a Kendrick, “Perspective: emerging evidence for signaling roles of mitochondrial anaplerotic products in insulin secretion.,” Am. J. Physiol. Endocrinol. Metab., vol. 288, no. 1, pp. E1–15, Jan. 2005.

[43] M. J. MacDonald et al., “Differences between human and rodent pancreatic islets: Low pyruvate carboxylase, ATP citrate lyase, and pyruvate carboxylation and high glucose-stimulated acetoacetate in human pancreatic islets,” J. Biol. Chem., vol. 286, no. 21, pp. 18383–18396, May 2011.

[44] R. Arrojo e Drigo, B. Roy, and P. E. MacDonald, “Molecular and functional profiling of human islets: from heterogeneity to human phenotypes,” Diabetologia, vol. 63, no. 10, pp. 2095–2101, 2020.

[45] J. N. Bazil, G. T. Buzzard, and A. E. Rundell, “Modeling mitochondrial bioenergetics with integrated volume dynamics.,” PLoS Comput. Biol., vol. 6, no. 1, p. e1000632, Jan. 2010.

[46] M. Huang, S. Paglialunga, J. M. K. Wong, M. Hoang, R. Pillai, and J. W. Joseph, “Role of prolyl hydroxylase domain proteins in the regulation of insulin secretion,” Physiol. Rep., vol. 4, no. 5, pp. 1–15, 2016.

[47] M. Hoang and J. W. Joseph, “The role of α-ketoglutarate and the hypoxia sensing pathway in the regulation of pancreatic β-cell function,” Islets, vol. 12, no. 5, pp. 108–119, 2020.

## References

[1] P. Detimary, J. C. Jonas, and J. C. Henquin, “Possible links between glucose-induced changes in the energy state of pancreatic b cells and insulin release. unmasking by decreasing a stable pool of adenine nucleotides in mouse islets.” The Journal of Clinical Investigation, vol. 96, no. 4, pp. 1738–1745, 10 1995. [Online]. Available: http://www.jci.org/articles/view/11s219

[2] P. Detimary, S. Dejonghe, Z. Ling, D. Pipeleers, F. Schuit, and J.-C. Henquin, “The changes in adenine nucleotides measured in glucose-stimulated rodent islets occur in β cells but not in α cells and are also observed in human islets,” Journal of Biological Chemistry, vol. 273, no. 51, pp. 33 905–33 908, 1998. [Online]. Available: http://www.jbc.org/content/273/51/33905.abstract

[3] P. Detimary, G. Van den Berghe, and J.-C. Henquin, “Concentration dependence and time course of the effects of glucose on adenine and guanine nucleotides in mouse pancreatic islets,” Journal of Biological Chemistry, vol. 271, no. 34, pp. 20 559–20 565, 1996. [Online]. Available: http://www.jbc.org/content/271/34/20559.abstract

[4] F. Wu, F. Yang, K. C. Vinnakota, and D. a. Beard, “Computer modeling of mitochondrial tricarboxylic acid cycle, oxidative phosphorylation, metabolite transport, and electrophysiology,” The Journal of biological chemistry, vol. 282, no. 34, pp. 24 525–37, 2007. [Online]. Available: http://www.ncbi.nlm.nih.gov/pubmed/175917s5

[5] S. M. Ronnebaum, O. Ilkayeva, S. C. Burgess, J. W. Joseph, D. Lu, R. D. Stevens, T. C. Becker, A. D. Sherry, C. B. Newgard, and M. V. Jensen, “A Pyruvate Cycling Pathway Involving Cytosolic NADP-dependent Isocitrate Dehydrogenase Regulates Glucose-stimulated Insulin Secretion,” Journal of Biological Chemistry, vol. 281, no. 41, pp. 30 593–30 602, 2006.

[6] I. R. Sweet and F. M. Matschinsky, “Mathematical model of beta-cell glucose metabolism and insulin release. I. Glucokinase as glucosensor hypothesis,” Am J Physiol Endocrinol Metab, vol. 268, no. 4, pp. E775–788, 1995. [Online]. Available: http://ajpendo.physiology.org/cgi/content/abstract/26s/4/E775

[7] N. Jiang, R. Cox, and J. Hancock, “A kinetic core model of the glucose-stimulated insulin secretion network of pancreatic *β*-cells,” Mammalian Genome, vol. 18, pp. 508–520, 2007, 10.1007/s00335-007-9011-y. [Online]. Available: http://dx.doi.org/10.1007/s00335-007-9011-y

[8] U. Wittig, M. Golebiewski, R. Kania, O. Krebs, S. Mir, A. Weidemann, S. Anstein, J. Saric, and I. Rojas, “Sabio-rk: Integration and curation of reaction kinetics data,” in Data Integration in the Life Sciences, ser. Lecture Notes in Computer Science, U. Leser, F. Naumann, and B. Eckman, Eds. Springer Berlin Heidelberg, 2006, vol. 4075, pp. 94–103.

[9] K. Yugi and M. Tomita, “A general computational model of mitochondrial metabolism in a whole organelle scale,” Bioinformatics, vol. 20, no. 11, pp. 1795–1796, 2004. [Online]. Available: http://bioinformatics.oxfordjournals.org/content/20/11/1795.abstract

[10] P. O. Westermark, J. H. Kotaleski, A. Bjorklund, V. Grill, and A. Lansner, “A mathematical model of the mitochondrial NADH shuttles and anaplerosis in the pancreatic beta-cell,” Am J Physiol Endocrinol Metab, vol. 292, no. 2, pp. E373–393, 2007. [Online]. Available: http://ajpendo.physiology.org/cgi/content/abstract/292/2/E373

[11] M. H. N. Hoefnagel, M. J. C. Starrenburg, D. E. Martens, J. Hugenholtz, M. Kleerebezem, I. I. Van Swam, R. Bongers, H. V. Westerhoff, and J. L. Snoep, “Metabolic engineering of lactic acid bacteria, the combined approach: kinetic modelling, metabolic control and experimental analysis.” Microbiology (Reading, England), vol. 148, no. Pt 4, pp. 1003–13, April 2002. [Online]. Available: http://www.ncbi.nlm.nih.gov/pubmed/11932446

[12] M. Scheer, A. Grote, A. Chang, I. Schomburg, C. Munaretto, M. Rother, C. Söhngen, M. Stelzer, J. Thiele, and D. Schomburg, “Brenda, the enzyme information system in 2011,” Nucleic Acids Research, vol. 39, no. suppl 1, pp. D670–D676, 2011. [Online]. Available: http://nar.oxfordjournals.org/content/39/suppl_1/D670.abstract

[13] U. Wittig, R. Kania, M. Golebiewski, M. Rey, L. Shi, L. Jong, E. Algaa, A. Weidemann, H. Sauer-Danzwith, S. Mir, O. Krebs, M. Bittkowski, E. Wetsch, I. Rojas, and W. Muller, “Sabio-rk-database for biochemical reaction kinetics,” Nucleic Acids Research, vol. 40, no. D1, pp. D790–D796, 2012. [Online]. Available: http://nar.oxfordjournals.org/content/40/D1/D790.abstract

[14] H. Schmidt and M. Jirstrand, “Systems biology toolbox for MATLAB^®^: a computational platform for research in systems biology,” Bioinformatics, vol. 22, no. 4, pp. 514–515, February 2006, http://www.sbtoolbox2.org.

[15] H. Schmidt, “https://iqmtools.intiquan.com/,” 2021. [Online]. Available: https://iqmtools.intiquan.com/

[16] D. Lu, H. Mulder, P. Zhao, S. C. Burgess, M. V. Jensen, S. Kamzolova, C. B. Newgard, and a. D. Sherry, “^13^C NMR isotopomer analysis reveals a connection between pyruvate cycling and glucose-stimulated insulin secretion (GSIS).” Proceedings of the National Academy of Sciences of the United States of America, vol. 99, no. 5, pp. 2708–13, March 2002. [Online]. Available: http://www.ncbi.nlm.nih.gov/pubmed/11ss0625

[17] A. Saltelli, Global Sensitivity Analysis: The Primer. John Wiley, Mar. 2008.

[18] A. C. Hindmarsh, P. N. Brown, K. E. Grant, S. L. Lee, R. Serban, D. E. Shumaker, and C. S. Woodward, “Sundials: Suite of nonlinear and differential/algebraic equation solvers,” ACM Trans. Math. Softw., vol. 31, no. 3, pp. 363–396, Sep. 2005. [Online]. Available: http://doi.acm.org/10.1145/10s9014.10s9020

[19] R. S. Malik-Sheriff, M. Glont, T. V. N. Nguyen, K. Tiwari, M. G. Roberts, A. Xavier, M. T. Vu, J. Men, M. Maire, S. Kananathan, E. L. Fairbanks, J. P. Meyer, C. Arankalle, T. M. Varusai, V. Knight-Schrijver, L. Li, C. Dueiias-Roca, G. Dass, S. M. Keating, Y. M. Park, N. Buso, N. Rodriguez, M. Hucka, and H. Hermjakob, “BioModels-15 years of sharing computational models in life science,” Nucleic Acids Research, vol. 48, no. D1, pp. D407–D415, 11 2019. [Online]. Available: https://doi.org/10.1093/nar/gkz1055

[20] K. Tiwari, S. Kananathan, M. G. Roberts, J. P. Meyer, M. U. Sharif Shohan, A. Xavier, M. Maire, A. Zyoud, J. Men, S. Ng, T. V. N. Nguyen, M. Glont, H. Hermjakob, and R. S. Malik-Sheriff, “Reproducibility in systems biology modelling,” Molecular Systems Biology, vol. 17, no. 2, p. e9982, 2021. [Online]. Available: https://www.embopress.org/doi/abs/10.15252/msb.202099s2

[21] P. Roger D, “Reproducible Research in Computational Science,” vol. 1226, no. 2011, 2014.

[22] W. S. Noble, “A quick guide to organizing computational biology projects.” PLoS computational biology, vol. 5, no. 7, p. e1000424, jul 2009. [Online]. Available: http://www.ncbi.nlm.nih.gov/pubmed/19649301

[23] S. M. Keating, D. Waltemath, M. König, F. Zhang, A. Dräger, C. Chaouiya, F. T. Bergmann, A. Finney, C. S. Gillespie, T. Helikar, S. Hoops, R. S. Malik-Sheriff, S. L. Moodie, I. I. Moraru, C. J. Myers, A. Naldi, B. G. Olivier, S. Sahle, J. C. Schaff, L. P. Smith, M. J. Swat, D. Thieffry, L. Watanabe, D. J. Wilkinson, M. L. Blinov, K. Begley, J. R. Faeder, H. F. Gómez, T. M. Hamm, Y. Inagaki, W. Liebermeister, A. L. Lister, D. Lucio, E. Mjolsness, C. J. Proctor, K. Raman, N. Rodriguez, C. A. Shaffer, B. E. Shapiro, J. Stelling, N. Swainston, N. Tanimura, J. Wagner, M. Meier-Schellersheim, H. M. Sauro, B. Palsson, H. Bolouri, H. Kitano, A. Funahashi, H. Hermjakob, J. C. Doyle, M. Hucka, and S. L. . C. members, “Sbml level 3: an extensible format for the exchange and reuse of biological models,” Molecular Systems Biology, vol. 16, no. 8, p. e9110, 2020. [Online]. Available: https://www.embopress.org/doi/abs/10.15252/msb.20199110

